# *Panx1* channels promote both anti- and pro-seizure-like activities in the zebrafish via *p2rx7* receptors and ATP signaling

**DOI:** 10.1101/2021.06.03.446992

**Authors:** Paige Whyte-Fagundes, Daria Taskina, Nickie Safarian, Christiane Zoidl, Peter L Carlen, Logan W Donaldson, Georg R Zoidl

## Abstract

The molecular mechanisms of excitation-inhibition imbalances promoting seizure generation in epilepsy patients are not fully understood. Experimental evidence suggests that Pannexin1 (*Panx1*), an ATP release channel, modulates excitability of the brain. In this report, we have performed behavioral and molecular phenotyping experiments on zebrafish larvae bearing genetic or pharmacological knockouts of *panx1a* or *panx1b* channels, each highly homologous to human PANX1. When Panx1a function is lost or both channels are under pharmacological blockage, treatment with pentylenetetrazol to induce seizures causes reduced ictal-like events and seizure-like locomotion. These observations were extended by transcriptome profiling, where a spectrum of distinct metabolic and cell signaling states correlates with the loss of *panx1a*. The pro- and anticonvulsant activities of both Panx1 channels affects ATP release and involves the purinergic receptor *p2rx7*. We propose that Panx1 zebrafish models offer opportunities to explore the molecular and physiological basis of seizures and assist anticonvulsant drug discovery.

## Introduction

A widely accepted view in epilepsy research is that neuronal hyperexcitability during epileptic seizures is caused by an imbalance of excitatory and inhibitory activities. This view places the imbalance of neuronal transmitters glutamate and GABA release first, but evidence is accumulating for altered ATP- and adenosine-mediated signaling between neurons and glial cells contributing to heightened states of excitability and epileptic seizures ^1^. ATP release and signaling can be both excitatory and inhibitory, but adenosine strongly inhibits electrical activity. Five groups of ATP-release channels with expression in the nervous system are known: pannexin-1 (Panx1) ^2^, connexin (Cx) hemichannels ^3^, calcium homeostasis modulator 1 (CALHM1) ^4^, volume-regulated anion channels (VRACs, also known as volume-sensitive outwardly rectifying (VSOR) anion channels) ^5^, and maxi-anion channels (MACs) ^6^. Panx1 is recognized as a pro-convulsant channel after behavioral and electrophysiological markers of excitability are ameliorated in distinct models of epilepsy once Panx1 is inhibited pharmacologically or by global deletion in mice ^7–10^. Other evidence for pro-convulsant actions of Panx1 derived from increased expression of human and rodent Panx1 found in epileptic tissue ^8,11–13^. However, the simplistic view of inhibiting mammalian Panx1 and causing anti-convulsant effects is challenged. The targeted deletion of mouse Panx1 in astrocytes potentiates, while the absence of Panx1 in neurons attenuates seizure manifestation ^14^. Furthermore, the contribution of Panx1 to seizures is also brain region dependent ^15^, raising questions about the underlying molecular and cellular mechanisms.

Zebrafish have two *panx1* ohnologues, *panx1a* and *panx1b,* with distinct expression localizations and biophysical properties ^16–18^. Like mammalian Panx1 ^19–21^, the zebrafish *panx1a* is broadly expressed in all tissues tested ^16,17^, whereas the expression of *panx1b* is highest in the nervous system ^17^. We had suggested that both pannexins fulfill different functions *in vivo* based on differences in the unitary conductance of Panx1a (≈380pS) and Panx1b (480-500pS) channels, the complexity of subconductance stages, and the cumulative open and closed times ^17^. Here, we interrogated Panx1 channels genetically and pharmacologically, and induced seizure-like activities in the zebrafish using pentylenetetrazole (PTZ) ^22^ to suppress inhibitory networks by blocking gamma aminobutyric acid (GABA)-A receptors. The zebrafish responses to PTZ were physiologically and behaviorally comparable to mammals ^22,23^.

Gene-edited zebrafish lacking *panx1a* or *panx1b* ^24,25^ revealed opposite seizure-like phenotypes and PTZ-related morbidity. Targeted ablation of *panx1b* potentiates, while the absence of *panx1a* attenuates seizure manifestations according to recordings of *in vivo* local field potentials (ivLFP) and locomotor behavior. A deletion of both fish *panx1* genes, in a double knockout fish (DKO), causes a moderate phenotype with reduced seizures. In line with these observations are significant changes to extracellular ATP levels and biological processes demonstrating that the propensity of developing seizure like activities is correlated with altered regulation of energy metabolism, cell death, and the cellular transportome. The acute pharmacological blocking of both Panx1 channels using Probenecid (Pb) abolished PTZ-induced seizures. The molecular, electrophysiological, and behavioral changes are overlapping, but not identical to genetic interference. Likewise, pharmacological blocking of the Panx1 interaction partner P2rx7 ^26^ reduces seizure-like activities like Pb treatment. Finally, structural modeling and comparing the pores of the two Panx1 ohnologues and the human PANX1 channel supports our experimental data implicating the Panx1a protein as a driver of pro-convulsant activities.

In summary, our analysis of genetic and pharmacological models in the zebrafish establishes Panx1a channels as the pro-convulsant Panx1 channel. The different propensities of Panx1a and Panx1b channels in developing seizure-like activities opens opportunities for comparative studies into seizure mechanisms and drug discovery by targeting shared properties of zebrafish Panx1a and human PANX1.

## Results

### Panx1 genotypes determine evoked seizure-like events

Seven days post fertilization (7dpf) larvae were anesthetized, and agar embedded before an electrode was placed into the optic tectum (OT). Seizure-like events (SLE) were recorded after topical application of 15 mM PTZ, as described ^27^ (**Fig. 1a).** Characteristic SLEs induced by PTZ demonstrate an initial action potential burst followed by short paroxysmal bursts, representative of a seizure pattern, which was absent at baseline (**Fig. 1b**). The number of *panx1a^−/−^* larvae exhibiting SLEs was reduced by 88% compared to PTZ treated Tubingen Longfin (TL) controls; only one *panx1a^−/−^* larva responded to PTZ treatment (n = 8). All *panx1b^−/−^* larvae exhibited SLEs (n = 9). 33% of the DKO larvae seized (*panx1a^−/−^: panx1b^−/−^;* DKO; n = 12), which was significantly reduced compared to controls (n = 7) and indistinguishable from *panx1a^−/−^* larvae (p=0.2) (**Fig. 1c**). The average number of SLEs per hour was similar for DKOs and *panx1a*^−/−^ larvae (p = 0.58), with less than one event on average (DKO: 0.58 ± 0.29; *panx1a^−/−^*: 0.38 ± 0.38 events/hour,). Even though all *panx1b^−/−^* fish had SLEs, the average number of events per hour were significantly less compared to controls (*panx1b^−/−^:* 36.78 ± 2.67; TL: 49.86 ± 3.86 events/hour) (**Fig. 1d**). The duration of events in all genotypes were comparable to TL controls (TL: 5.5s ± 0.4s; DKO: 4.1s ± 0.2s, p = 0.07; *panx1a*^−/−^: 4.2s, p = n/a; *panx1b^−/−^*: 4.8s ± 0.4s, p = 0.09) (**Fig. 1e**).

**Fig. 1.**
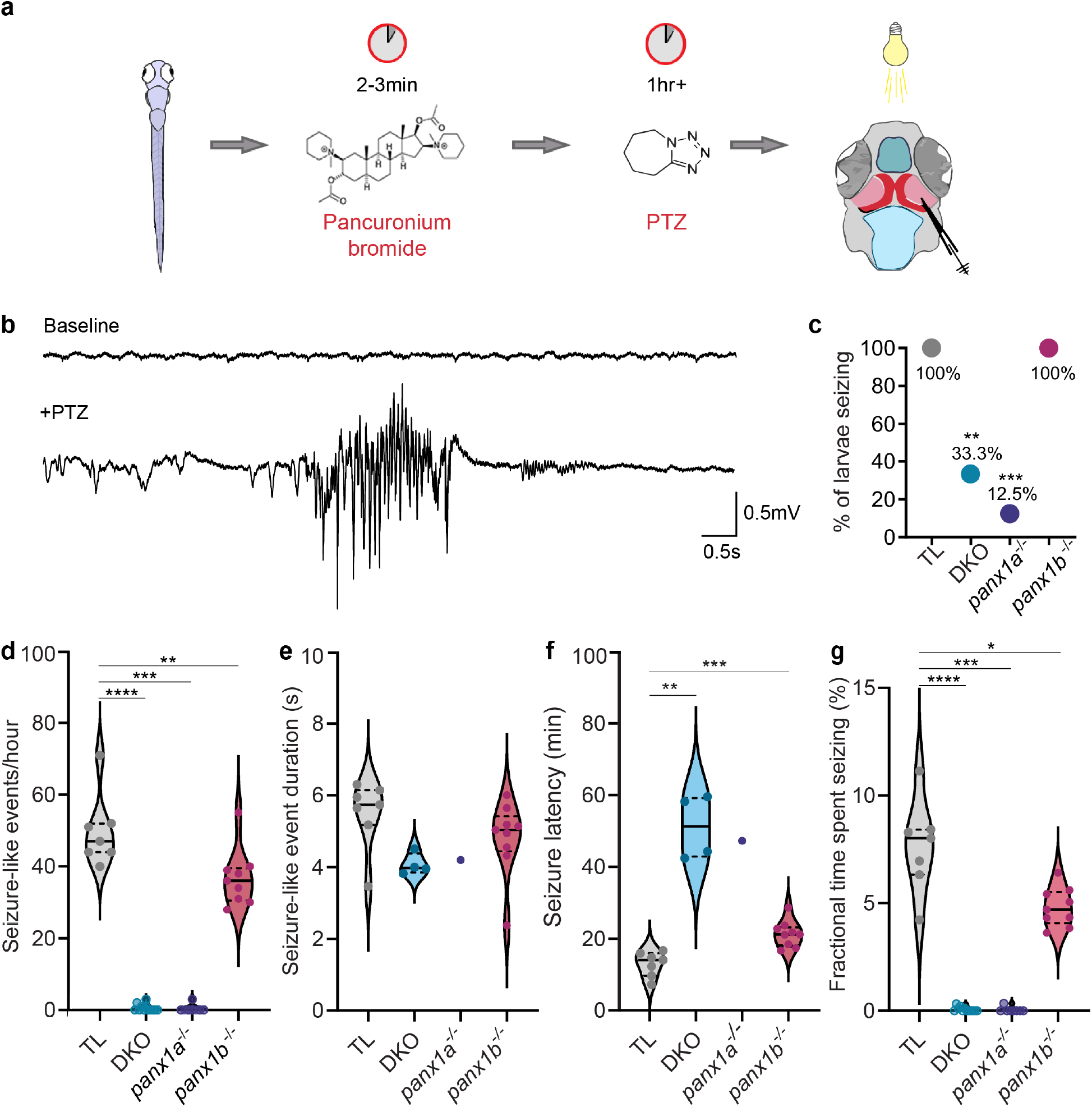
*Panx1* knockout larvae showed distinct seizure-like activities *in vivo*. **a** Workflow for recording *in vivo* local field potentials (ivLFP) from the right optic tectum of 7dpf anesthetized larva after 15mM PTZ treatment. **b** Representative recording of baseline activity (top) from TL larvae and a seizure-like event (SLE) sample induced with the addition of PTZ. **c** All TL and *panx1b^−/−^* larvae had SLEs (TL (grey) n = 7/7; DKO (light blue) n = 4/12; *panx1a^−/−^* (deep blue) n = 1/8; *panx1b^−/−^* (magenta) n = 9/9). **d** Quantification of SLEs in the first hour of ivLFP recording revealed that all *panx1* knockout (KO) larvae had a significant reduction in SLEs compared to PTZ treated TL controls, presented as an average number of SLEs/hour ± s.e.m. **e** The average duration of SLEs (in seconds ± s.e.m.) for each genotype were not significantly different compared to TLs. Statistical tests were not significant for *panx1a^−/−^* due to lack of statistical power. **f** All *panx1* knockout larvae had a significant delay in the average onset time (minutes ± s.e.m.) of the first seizure-like event compared to PTZ treated TL controls. **g** All *panx1* knockout larvae spent significantly less time seizing compared to PTZ treated TL controls. Average fractional time spent seizing is presented in percent ± s.e.m. Scale bars 0.5mV by 0.5s. *p < 0.05, **p < 0.01, ***p < 0.001, ****p < 0.0001.

The latency until the first SLE showed differences amongst genotypes. TL larvae exhibited the shortest latent period (12.9 ± 1.3min). *Panx1b^−/−^* fish presented a delay until the first SLE, with events starting at 21 ± 1.2min. DKO (51.1 ± 4.5min) and *panx1a*^−/−^ responded last to PTZ treatment (47.3min) (**Fig. 1f**).

The fractional percentage of seizing time within the first hour revealed additional genotype specific differences. SLEs in *panx1b^−/−^* larvae were reduced compared to TL controls (*panx1b ^−/−^*: 4.8% ± 0.3%; TL: 7.6% ± 0.8%). However, both DKO and *panx1a^−/−^* fish displayed minimal seizing time (<0.1%). (**Fig. 1g**).

### Loss of Panx1 caused local network differences in the optic tectum of PTZ treated larvae

The time-frequency domain was examined by visual inspection of ivLFPs combined with an in-house algorithm to automate SLE detection ^28,29^. Representative one-hour traces of electrographic recordings for TL controls, DKO, *panx1a^−/−^*, and *panx1b^−/−^* after PTZ application are shown in **Fig. 2a-d**. The exposure to PTZ elicited discharges characterized by large amplitudes and poly-spiking activities of at least 3 seconds in duration. TL controls and *panx1b^−/−^* larvae showed typical SLEs (red dotted lines zoom into these regions). Highlighted events near the end of the representative traces for DKO and *panx1a^−/−^* larvae demonstrated the lack of seizure-like electrographic signatures (blue dotted lines zoom into these regions). LFPs revealed changes in spectral power in the time and frequency domains; visible in expanded views of the ivLFPs and corresponding spectrograms. TL controls (**2a**) and *panx1b*^−/−^ (**2d**) demonstrated a robust increase in low-frequency power and increased power in high frequencies above 60Hz. The spectrograms for DKO (**2b**) and *panx1a^−/−^* (**2c**) were scaled to match TL (**2a**) and *panx1b^−/−^* (**2b**) data and revealed frequency power like baseline activity (**Supplementary Fig. 1**).

**Fig. 2.**
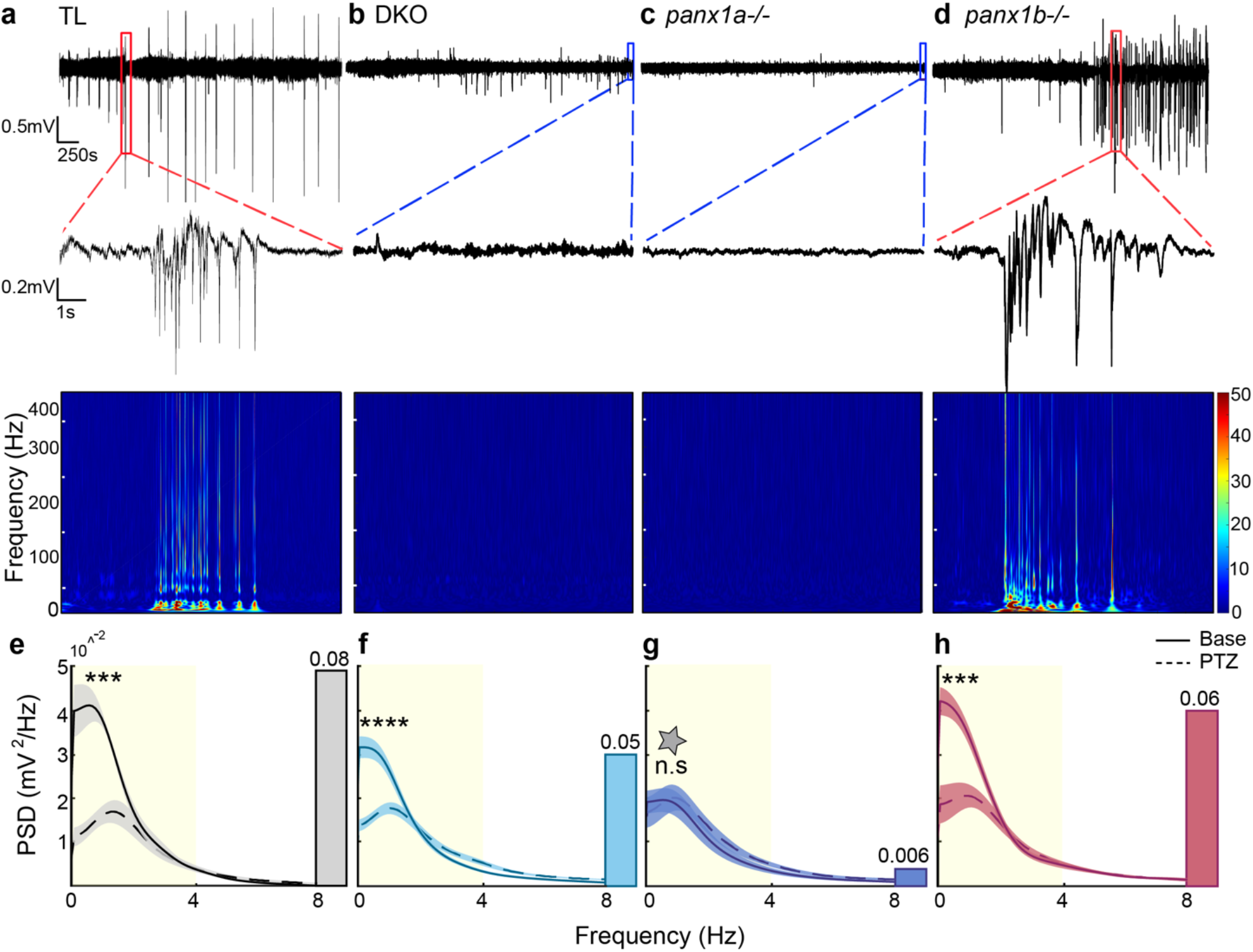
Spectral Analysis of LFPs revealed unique differences in larvae. Representative one-hour long LFP traces with PTZ treatment: **a** TL **b** DKO, **c** *panx1a^−/−^* and **d** *panx1b^−/−^*. TL and *panx1b^−/−^* traces showed typical seizure-like events. Single events were highlighted by red dotted lines and magnified. Blue dotted lines highlight lack of seizure-like activity in DKO and *panx1a^−/−^* larvae. Spectrograms below the traces showed increases in low and high frequency power during seizure-like events for TL and *panx1b^−/−^*. **e – h** Power spectral density (PSD, mV^2^/Hz) measured across frequencies revealed no significant differences in baseline (dotted lines) frequencies for all genotypes (p = 0.42; **e**: TL, grey; **f**: DKO, light blue; **g**: *panx1a^−/−^,* dark blue; **h**: *panx1b^−/−^,* magenta). The power was significantly increased in the delta band (1-4 Hz) after PTZ treatment (solid line) for all genotypes except *panx1a^−/−^*. The PSD was plotted as an average across traces with the shaded regions indicating ± s.e.m. Changes in delta power were quantified from the areas under the curves (AUC) of the power spectrum (in yellow). The changes in AUCs are represented as a bar graphs inserted to the right of the power spectra. TL showed the greatest change in delta, followed by *panx1b^−/−^*, and DKO. *Panx1a^−/−^* exhibited insignificant changes and maintained delta power throughout recordings. Scale bars: top = 0.5mV by 250s and bottom = 0.2mV by 1s. ***p < 0.001, ****p < 0.0001, grey star - p = 0.0015 compared to TL with PTZ.

Power spectral density was measured to identify tectal network differences. Baseline activities were similar across genotypes (p = 0.42 for each comparison). Changes were seen in delta power (highlighted in yellow) after PTZ treatment and quantified by measuring the difference in the area under the power spectrum from 1 – 4Hz (inset bar graphs) (**Fig. 2 e-h**). PTZ treatment significantly increased delta power for TL (**2e**), DKO (**2f),** and *panx1b^−/−^* larvae (**2h**). In contrast, delta power for *panx1a^−/−^* remained at base level (**2g**; dark blue bar). Delta power of *panx1a^−/−^* larvae during PTZ treatment was significantly decreased compared to PTZ treated TL controls (**2g;** grey star, p = 0.0015). *Panx1b^−/−^* (p = 0.99) and DKO larvae (p = 0.25) showed similar delta power during PTZ treatment compared to TL(TL = 0.08 (grey bar), *panx1b*^−/−^ = 0.06 (magenta bar), DKO = 0.05 (light blue bar)).

We concluded that the electrical discharges for both TL controls and *panx1b^−/−^* larvae were similar in waveform to those previously reported in zebrafish ^22,27^. However, they occurred less frequently in *panx1b^−/−^* fish and were absent in *panx1a^−/−^* larvae.

### *DKO and panx1a^−/−^ larvae* have reduced interictal-like activity in the optic tectum

Interictal-like epileptiform discharges (ILED) were investigated for genotype-dependent changes in occurrence and waveform. In ivLFP data, ILEDs were defined as events that were shorter than 3 seconds and had amplitudes greater than 1.5 times the baseline activity ^30^. **Fig. 3a** depicts typical interictal-like events, with TL (grey line) and *panx1b^−/−^* (magenta line) appearing most similar. The quantification of ILEDs during a one-hour PTZ application period showed that DKO (119 ± 7.2 events/hour) and *panx1a^−/−^* (121.4 ± 12.9 events/hour) larvae exhibited interictal-like events. However, their ILEDs occurred significantly less when compared to TL (167.3 ± 9.3 events/hour) and *panx1b^−/−^* larvae (174.1 ± 11.5 events/hour); both TL and *panx1b^−/−^* were similar.

**Fig. 3.**
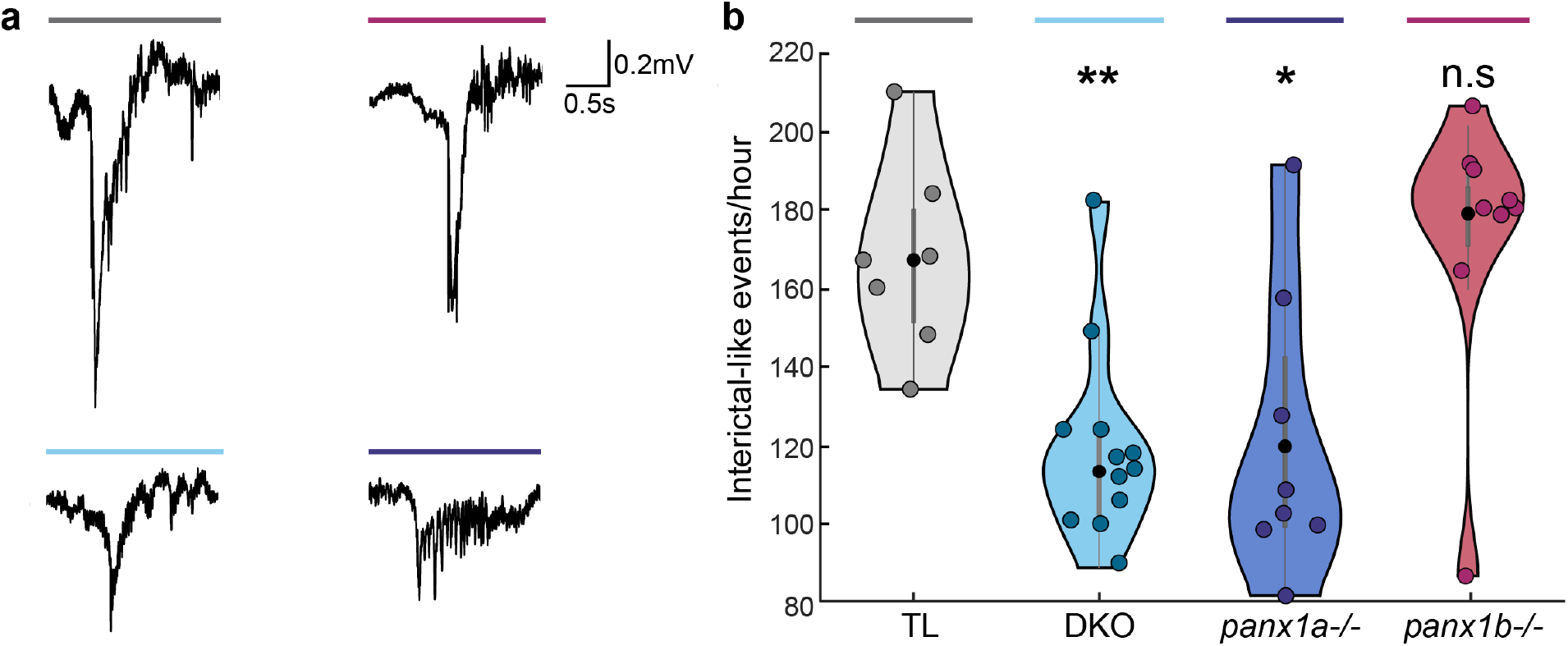
Interictal-like activity was decreased in the optic tectum of DKO and *panx1a^−/−^* larvae. **a** Electrophysiological recordings of PTZ treatment showed interictal-like epileptiform discharges (ILED) in all genotypes; representative ILEDs top: TL (grey line) and *panx1b^−/−^* (magenta line) shows most similarities and bottom: DKO (light blue line) and *panx1a^−/−^* (deep blue line). **b** Quantification of ILEDs for the first hour of recording for all genotypes revealed that DKO and *panx1a^−/−^* had significantly fewer ILEDs. No significant difference was found in the amount of ILEDs for *panx1b^−/−^* larvae compared to TL. Data presented as an average number of events per hour ± s.e.m. Scale bar: 0.2mV by 0.5s. *p < 0.05, **p < 0.01.

### Targeting Panx1 improves PTZ-induced seizure locomotion and molecular responses

Locomotion tracking was used to quantify genotype-specific seizure-related behaviors in response to 15 mM PTZ treatment (**Fig. 4a**). Activity scores (Δpixel) for locomotor activity were plotted for: rest, baseline, and post-PTZ treatment (**Fig. 4b**). At the start, addition of fresh E3 medium caused a transient minor activity increase for all larvae (n = 36 per genotype), which stabilized at baseline within 30min. PTZ treatment increased locomotor activity significantly in all genotypes. TL larvae showed a continuous activity increase in the first 10 minutes, a peak around 15 to 30min and a gradual decline within the hour. TL and DKO activity curves were statistically similar. The *panx1a^−/−^* activity curve was significantly reduced compared to TL controls after the transient peak observed 20min post-PTZ treatment. The *panx1b^−/−^* larvae exhibited a sharp spike in activity between 10 to 20min, with a steeper decline compared to TL.

**Fig. 4.**
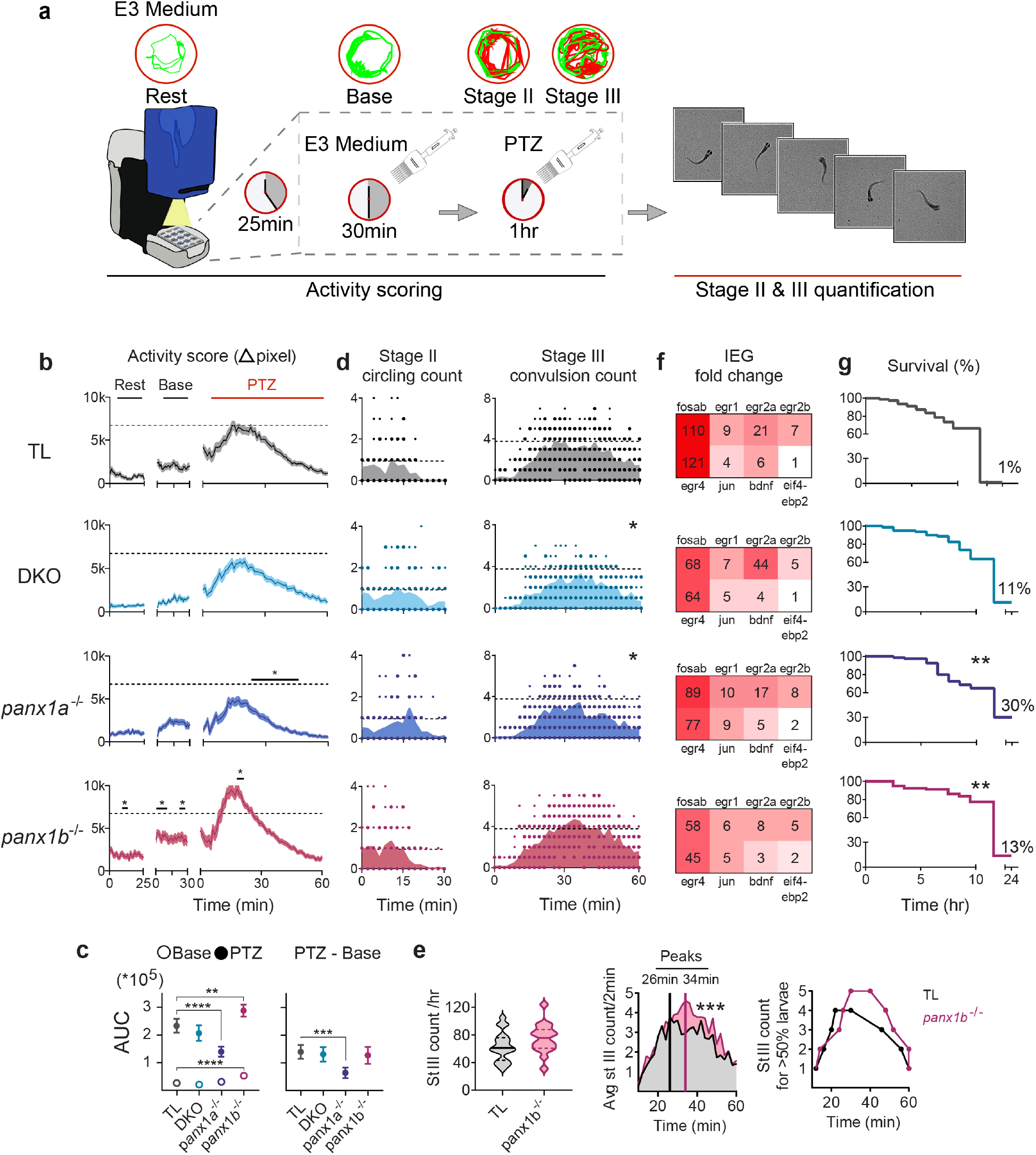
Behavioral and molecular outcomes of genetically targeting *panx1*. **a** Methodological workflow of behavioral assays in TL and *panx1* knockout larvae. **b** Larvae’s (n =36) baseline and 15mM PTZ-induced activity (Δpixel ± s.e.m.) were scored. *Panx1b^−/−^* baseline activity was higher than TL. PTZ induced hyperactivity peaked at 15-30 minutes for TL (grey, top), was reduced in DKO (blue, second), significantly reduced in *panx1a^−/−^* (blue, third) for the last 40 minutes, and significantly increased in *panx1b^−/−^* larvae (red, bottom) around 20min compared to TL. Dashed lines indicate TL’s max average activity score. **c** Left: AUC derived from the activity plots in **b** significantly differed between TL and *panx1b^−/−^* baselines (open points). AUC for one-hour post-PTZ treatment was significantly reduced for *panx1a^−/−^* and increased for *panx1b^−/−^* (filled points). Right: Change in PTZ treatment from average baseline activity was significantly reduced in *panx1a^−/−^* compared to TL. **d** Stage II count (rapid circling) did not differ among groups (n = 18; count/2min). Stage III (convulsion) was significantly reduced in DKO and *panx1a^−/−^* compared to TL. Dashed lines represent max average stage II and stage III counts for TL. **e** Left: Total stage III count between TL and *panx1b^−/−^* did not significantly differ. Middle: Time course of average stage III counts significantly differed between the two groups, peaking at 26min post-PTZ treatment for TL and 34min for *panx1b^−/−^*. Right: Time course where >50% of the larvae reached 1-5 stage III counts revealed a delayed peak onset for *panx1b^−/−^*. **f** IEG upregulation after one-hour PTZ treatment, represented as fold-change for PTZ treated larvae against non-treated controls. *Panx1* knockout larvae had reduced upregulation of the IEGs compared to TL, with DKO and *panx1b^−/−^* showing the greatest reduction. **g** Survival rate of larvae (n = 80) post PTZ treatment was significantly higher for *panx1a^−/−^* followed by *panx1b^−/−^,* and survival rate of DKOs did not differ from TL. Survival at 24 hours: TL: 1.25%, DKO: 11%, *panx1a^−/−^*: 30%, *panx1b^−/−^*: 13%. *p < 0.05, **p < 0.01, ***p < 0.001, ****p < 0.0001.

PTZ-induced hyperactivity was quantified from the area under the curve (AUC) shown in Figure 4b (**Fig. 4c**). The baseline AUC was significantly larger in *panx1b^−/−^ larvae* relative to TL in the last 15 minutes of stabilized baseline activity. The AUC post-PTZ treatment, taken over 60min, was greater in *panx1b^−/−^* and reduced in *panx1a^−/−^* relative to TL. When the mean baseline activity was extracted from the post-PTZ treatment AUC, only *panx1a^−/−^* activity was significantly reduced when compared to TL.

Consecutive stages of seizure-like behavior were analyzed next (**Supplementary Videos 1, 2**). Stage II, a rapid ‘whirlpool-like’ circular movement, and stage III, uncontrollable clonus-like twitching of the body followed by a prolonged loss of posture, were scored manually via video recordings of the fish. Stage II and III events were quantified in 2-minute intervals for one hour of PTZ treatment (n = 18 per genotype; **Fig. 4d**). Stage II onset occurred within a few minutes of treatment and lasted for 30min. TL and *panx1b^−/−^* entered Stage II first and peaked at ≈10 minutes post-treatment. Stage II activities for DKO and *panx1a^−/−^* occurred uniformly through-out the first 30min. The onset and total count of stage II locomotion did not significantly differ between genotypes. Stage III convulsive behavior started after 10min of PTZ treatment for all genotypes. Peak stage III activity was observed within 20 to 40min followed by a gradual decline. Stage III events in DKO and *panx1a^−/−^* were significantly reduced compared to TL controls. The total count of stage III events did not differ between *panx1b^−/−^* (74.9 ± 5 count/hour) and TL (63.2 ± 5 count/hour; **Fig. 4e left**). However, the peak activity latency, calculated at the time point with the highest average of stage III activity, was significantly delayed for *panx1b^−/−^* (34min) compared to TL (26min; **Fig. 4e middle**). Furthermore, time points at which 50% or more larvae entered stage III, indicated a delayed onset and termination of the stage III in *panx1b^−/−^* (**Fig. 4e right**).

Differential expression of Immediate Early Genes (IEGs) was tested after the behavioral assays’ experimental endpoint was reached. The IEGs were selected based on previously reported robust activation in a zebrafish PTZ model ^22^, in the human epileptic neocortex ^31^, and rodent models ^32,33^. **Fig. 4f** shows a 100-fold increase in the expression of *fosab* and *egr4* (in red) in PTZ treated TL, which was twice as high compared to the fold change observed in DKO and *panx1b^−/−^*. DKO and *panx1b^−/−^* larvae showed a modest upregulation for *fosab, egr1, egr2b, egr4, bdnf*. In *panx1a^−/−^* larvae, upregulation of *fosab, egr2a, egr4, bdnf* was moderate, but *egr1, egr2b,* and *jun* showed a strong differential upregulation **(Supplementary Table 1)**.

Kaplan-Meier plots show the larvae survival, which was determined by monitoring blood circulation and heartbeat post-PTZ treatment for ten consecutive hours and again at 24 hours (n = 80 per genotype; **Fig. 4g**). Although the p*anx1a*^−/−^ group displayed a decline between 5 to 10 hours, they had the highest survival rate of 30% at 24 hours. The *panx1b^−/−^* group’s survival rate was stable for the first 10 hours and declined to 13% after 24 hours. The low DKO and TL’s survival rate at 24 hours was statistically similar.

### Acute pharmacological blocking of panx1 suppresses seizure-like activities

Pharmacological blocking with Probenecid (Pb), a well-established Panx1 channel blocker ^34^, was used to determine acute changes to electrophysiological discharges, behavioral and molecular outcomes in the presence of PTZ. Valproic acid (VPA), a common anti-convulsant, was applied as a control for the suppression of SLEs ^35^. Electrodes were placed in the right optic tectum. All TL larvae were treated with Pb (75µM) or VPA (5mM) for 10min prior to PTZ application (**Fig. 5a**). For the duration of PTZ treatment, ivLFPs were monitored and confirmed SLE suppression in all Pb (n = 7) and VPA (n = 7) treated larvae (**Fig. 5b**). Sample recordings from larvae treated with PTZ and Pb (**Fig. 5c**) or VPA (**Fig. 5d**) showed no SLEs. Traces from timepoints near the experimental endpoint were depicted at higher resolution to show the larger deflections that occurred. The spectral analysis of these deflections revealed that no increase in high frequency power was associated with this type of activity. To confirm that Pb was not supressing SLEs due to toxicity, ivLFPs were recorded from larvae exposed to Pb or VPA without PTZ and compared to larvae with no drug treatments. Electrographic activity was comparable amongst these groups (**Fig. 5e, Supplementary Fig. 2**).

**Fig. 5.**
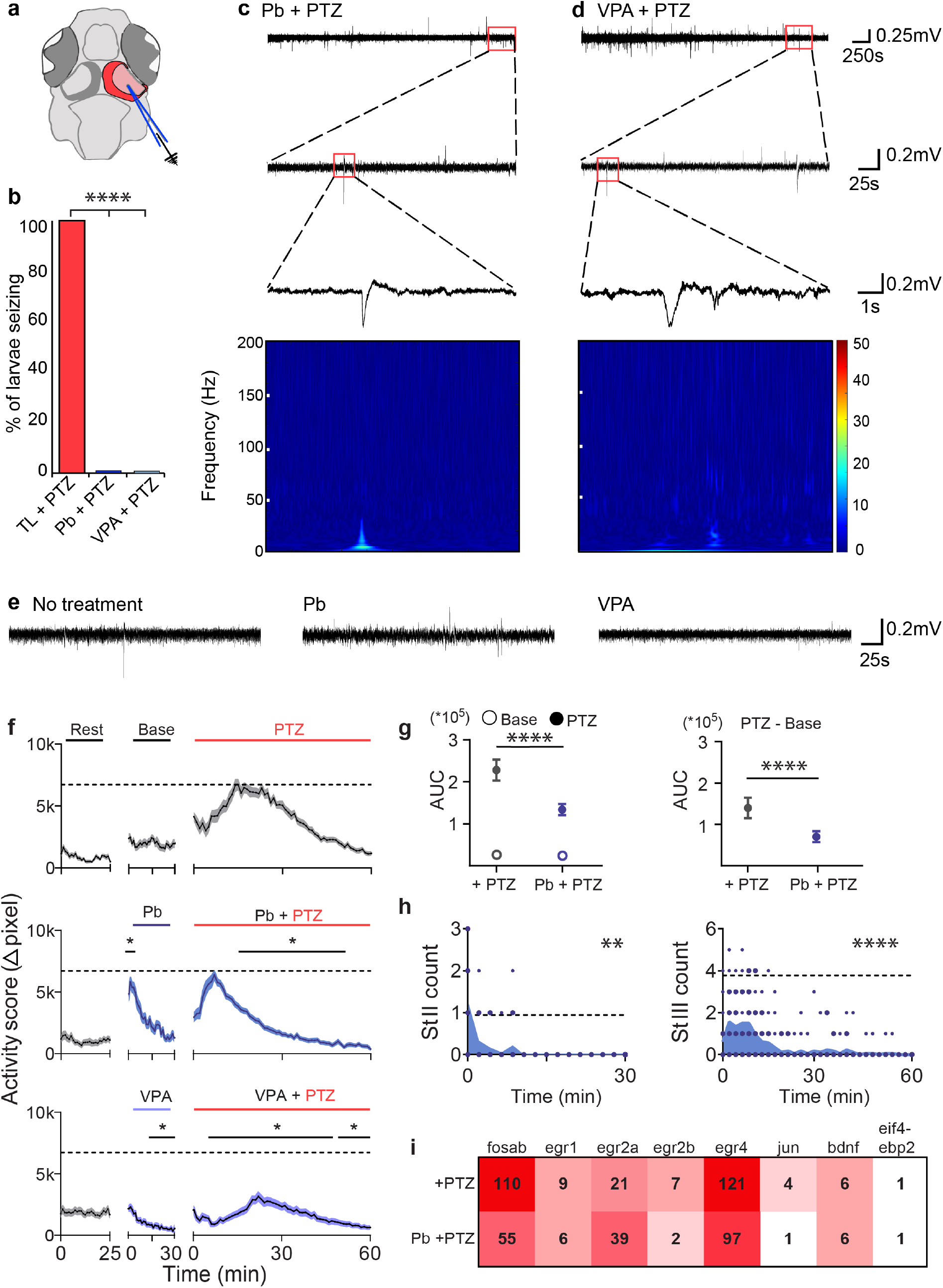
Acute blocking of Panx1 with Pb is effective for preventing seizure-like activity. **a** Schematic highlighting the region that LFPs were recorded from. **b** Larvae treated with 75µM Pb (n = 7) or 5mM VPA (n = 7) have no PTZ inducible SLEs compared to 100% of TL larvae (n = 7). Representative traces of larvae treated with **c** Pb and **d** VPA treated with PTZ for 60min. Expanded views into the last 5min of traces showed lack of SLEs. Spectrograms corresponding to small spikes in traces above demonstrate no increase in high frequency power associated with these events. **e** LFPs of the last 5min of 60min traces revealed no drug induced changes to baseline activity. Left: larvae with no treatment; middle: Pb treatment only; Right: VPA treatment only. **f** Baseline activity (Δpixel ± s.e.m.; n = 36) increased with Pb treatment for the first 5min. PTZ-induced hyperactivity subsided significantly within 15min, sooner than the PTZ only group. VPA treatment (n = 60) decreased baseline and PTZ-induced activity, with a hyperactivity curve like the PTZ only group. Dashed lines indicate max average activity for PTZ only group. **g** AUCs for baseline activity and Pb treatment did not differ (left, open points). Pb treatment significantly reduced the effect of PTZ without (left, filled points) and with (right) extracted baseline activity. **h** Stage II and III counts (n = 18; count/2min) were significantly reduced with Pb treatment, majority occurring in the first 10min of treatment. Dashed lines represent max average Stage II and Stage III counts for PTZ-only group. **i** IEG upregulation was reduced in Pb treated TL larvae except for *egr2a*. Scale bars: Top = 0.25mV by 250s, middle = 0.2mV by 25s, bottom = 0.2mV by 1s, **e** = 0.2mV by 25s. *p < 0.05, **p < 0.01, ***p < 0.001, ****p < 0.0001.

Locomotor activity was monitored for TL larvae treated with either 75µM Pb (n = 36) or 5mM VPA (n = 60) and compared to untreated controls (**Fig. 5f**). Pb application alone induced a brief, but sharp, increase in larvae activity that returned to baseline within 30min (**Fig. 5f middle**). VPA treatment caused reduced activity, which decreased below baseline after 10min (**Fig. 5f bottom**). After a 30min baseline recording, all larvae were treated with PTZ for 60min. Pb treated larvae had an early onset of spiking activity within the first 10min after PTZ treatment, ahead of larvae treated with PTZ only. Pb treated PTZ-induced activity subsided within 30min and returned to baseline, significantly faster than PTZ only treated larvae. VPA treatment significantly reduced PTZ-induced activity, with a similar timeline and curve profile to the PTZ-only group. AUC analysis on activity plots in **Fig. 5f** showed that baseline AUC did not significantly differ between no treatment and Pb treated larvae in the last 15min of stabilized activity (**Fig. 5g**). Post-PTZ treatment AUC, with or without extracted baseline activity, was larger in the PTZ-only treated group compared to the Pb-PTZ treated group. The analysis of seizure-related Stages II and III of Pb treated larvae (n = 18) every 2min for a duration of 60min (**Fig. 5h**) showed that Stage II activity was significantly reduced compared to PTZ-only treated larvae, with most of the activity occurring in the first 10min. Stage III activity of Pb treated larvae displayed an immediate onset compared to the PTZ-only group and was significantly reduced after 20min of PTZ treatment. The differential expression of IEGs after Pb treatment was consistent with the changes observed in gene-edited larvae; IEG expression was reduced after Pb treatment when compared to the PTZ only treated group (**Fig. 5i**). Pb treated TL larvae exhibited IEG expression like DKOs, where all IEGs exhibited less upregulation, except for *egr2a*. *Fosab’s* upregulation was reduced by 2-fold and *jun* had no significant upregulation with Pb treatment (**Supplementary Table 2**).

### Seizure phenotypes are linked to transcriptome changes

RNA-seq data representing the transcriptomes of untreated 6dpf larvae ^24,36^ were analyzed next. Venn diagrams highlighted considerable differences in the number and regulation of genes when the FishEnrichR database ^37,38^ was data-mined for biological processes enriched in the nervous system **(Supplementary Fig. 3)**. The GO biological process categories selected for further analysis represented broad themes based on known (signal transduction, cell death, transport) or anticipated (metabolism, cellular respiration) roles of *Panx1* channels (**Fig. 6a, Supplementary Table 3**).

**Fig. 6.**
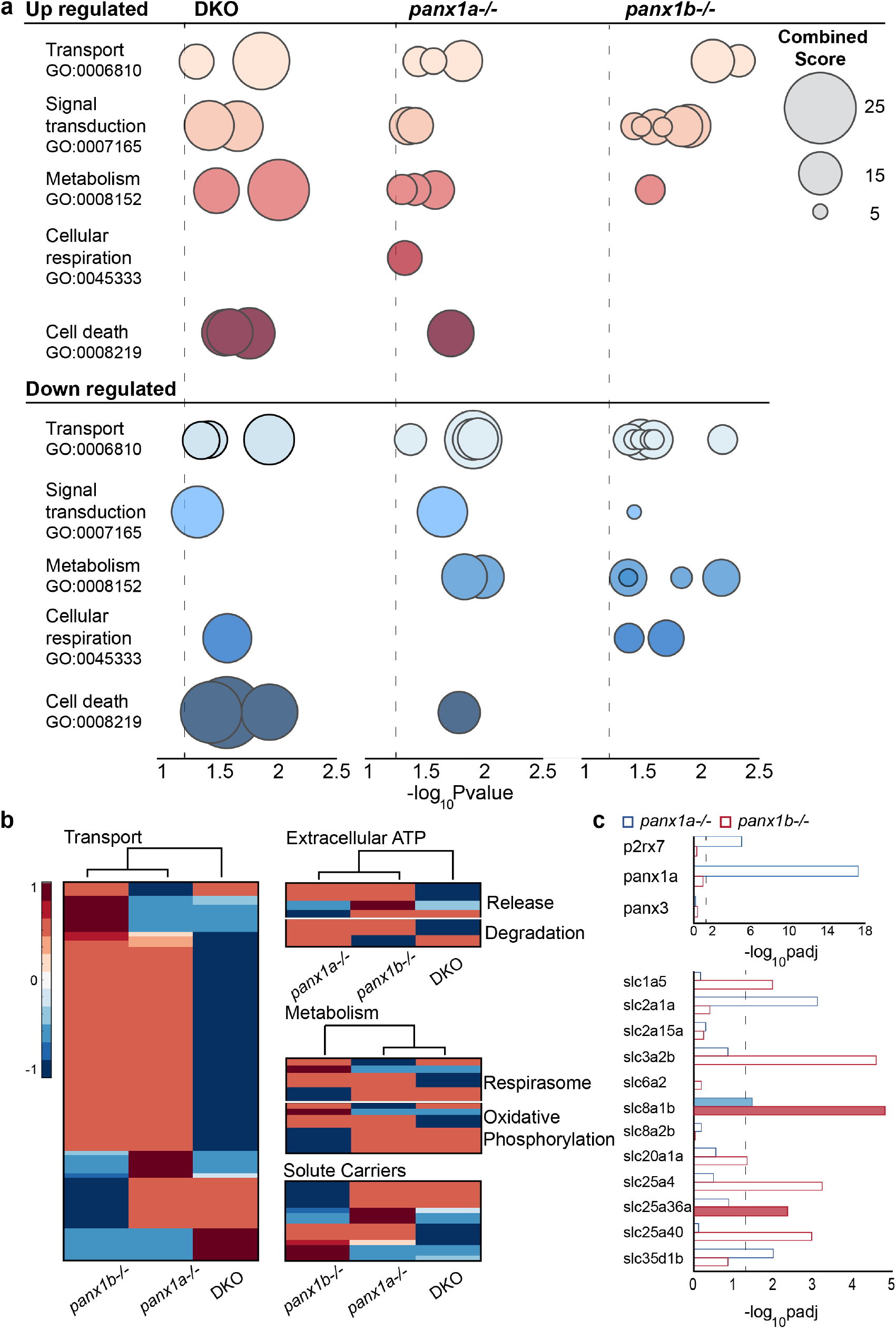
Loss of *Panx1a* affects biological processes related to metabolism and transport. **a** Gene ontology enrichment of biological processes for differentially regulated genes from RNA-seq data of DKO (left), *panx1a^−/−^* (middle) and *panx1b^−/−^* (right) found using the FishEnrichR database. Biological processes were grouped into 5 categories; transport (GO:0006810), signal transduction (GO:0007165), metabolism (GO:0008152), cellular respiration (GO:0045333) and cell death (GO:0008219). Data is presented as -log *p*-value based upon the fisher exact test and dot size represents the combined score for genes associated with that pathway. Dotted grey lines indicate *p*-value = 0.05. **b** Candidate genes were selected and RNA-seq data was mined for differential regulation in panx1 fish lines. Clustergrams compare *panx1* knockout larvae for correlation amongst chosen genes for (1) transport, (2) extracellular ATP (release & degradation), (3) metabolism (respirasome & oxidative phosphorylation) and (4) solute carriers. Scale shows correlation: red is positive, and blue is negative. Hierarchical cluster branches are shown above each clustergram and labelled below. **c** Expression (logpadj) of genes part of the ATP release (top) and solute carrier (bottom) clustergrams for *panx1a^−/−^* (blue) and *panx1b^−/−^* (top). Filled bars indicate upregulated genes, empty bars are downregulated. *p*-value = 0.05 indicated with dotted line.

All genotypes shared enrichment of the significant categories, *transport* (GO:0006810) and *signal transduction* (GO:0007165). The lack of enrichment of *cellular respiration* (GO:0045333) in *panx1b^−/−^* larvae suggested that mitochondrial energy metabolism processes and response to reactive oxygen species were less controlled after losing *panx1b*. Further, *panx1b^−/−^* larvae stood out for lacking enrichment of *cell death-associated processes* (GO:0008219). The opposite enrichment of the same categories in *panx1a^−/−^* larvae repeated the pattern of electrographic and behavioral activities. DKOs displayed a mixed phenotype, lacking a downregulated metabolism (GO:0008152).

Cluster analysis was performed to link coordinated expression changes with *Panx1* genotypes (**Fig 6b**). Genes for cluster analysis were manually curated based on assumptions made from the enriched biological processes (**Supplementary Table 4)**. Genotype-specific transport and metabolic properties corroborated the pattern of shared and distinct roles of zebrafish *panx1* ohnologs. *Panx1a^−/−^* and *panx1b^−/−^* larvae had the opposite coordination of gene expression changes in subcategories of metabolism; mitochondrial ATP production and oxidative phosphorylation processes. However, the coordinated expression was strengthened for genes encoding proteins that release or degrade ATP in *panx1a^−/−^* and *panx1b^−/−^* larvae. Interestingly, transport genes showed similar enhanced coordination in both genotypes consistent with roles of both channel proteins in cell signaling. However, coordinated expression of the solute carrier (SLC) group of membrane transport proteins involved in signal transduction pathways showed more significant variation between genotypes consistent with the GO ontology enrichment analysis; overall, DKOs had a less coordinated expression in these categories.

A direct comparison of *panx1a^−/−^* and *panx1b^−/−^* larvae corroborated the significant downregulation of *panx1a* and *p2rx7* expression as a major difference between the two genotypes (**Fig 6c**). Interestingly, the *slc2a1a* was downregulated in *panx1a^−/−^* larvae. In mammalia this glucose transporter is in the blood-brain barrier. Pathogenic SLC2A1 variants are associated with epilepsy ^39^. In *panx1b*^−/−^ larvae, SLC proteins with molecular functions in Na^+^/Ca^2+^ transmembrane transport (*slc8a1b*), or as pyrimidine nucleotide transmembrane transporters (*slc25a36a*) were upregulated, while regulation of sodium dependent amino acid transporters (*slc1a5*), mitochondrial ADP/ATP antiporters (*slc25a4*), or transport across the inner membrane of mitochondria (*slc25a40*), was reduced. Although the exact function of the upregulated *slc3a2b* in the zebrafish is unknown, in humans, heterodimers of SLC3A2 with the large neutral amino acid transporter (LAT1) function as sodium-independent, high-affinity transporters that mediate uptake of large neutral amino acids such as phenylalanine, tyrosine, L-DOPA, leucine, histidine, methionine, and tryptophan. Although the analysis presented only a snapshot of relevant molecular signatures, we concluded that the differences in prominent biological processes primed each genotype differently for the treatment with PTZ.

### Evidence for an ATP-dependent mechanism contributing to seizure-like activity

Changes to ATP release and differential expression of biomarkers in response to PTZ conditions were quantified as shown in **Fig. 7a**. Extracellular ATP concentrations [ATP]_ex_ were normalized to TL controls (grey bars) with and without PTZ treatment (baseline). Baseline [ATP]_ex_ (blue bars) was significantly reduced for *panx1a^−/−^* and Pb-treated TL larvae (**Fig. 7b**). PTZ treatment (red bars) did not change [ATP]_ex_ for *panx1a^−/−^* and Pb-treated TL larvae, which was consistent with reduced seizure-like activities. In DKO and *panx1b^−/−^* larvae, [ATP]_ex_ was significantly elevated overall compared to TL controls and was substantially altered by PTZ treatment. Our results suggested that the propensity of developing SLEs was correlated with regulation of energy metabolism and the availability of extracellular ATP.

**Fig. 7.**
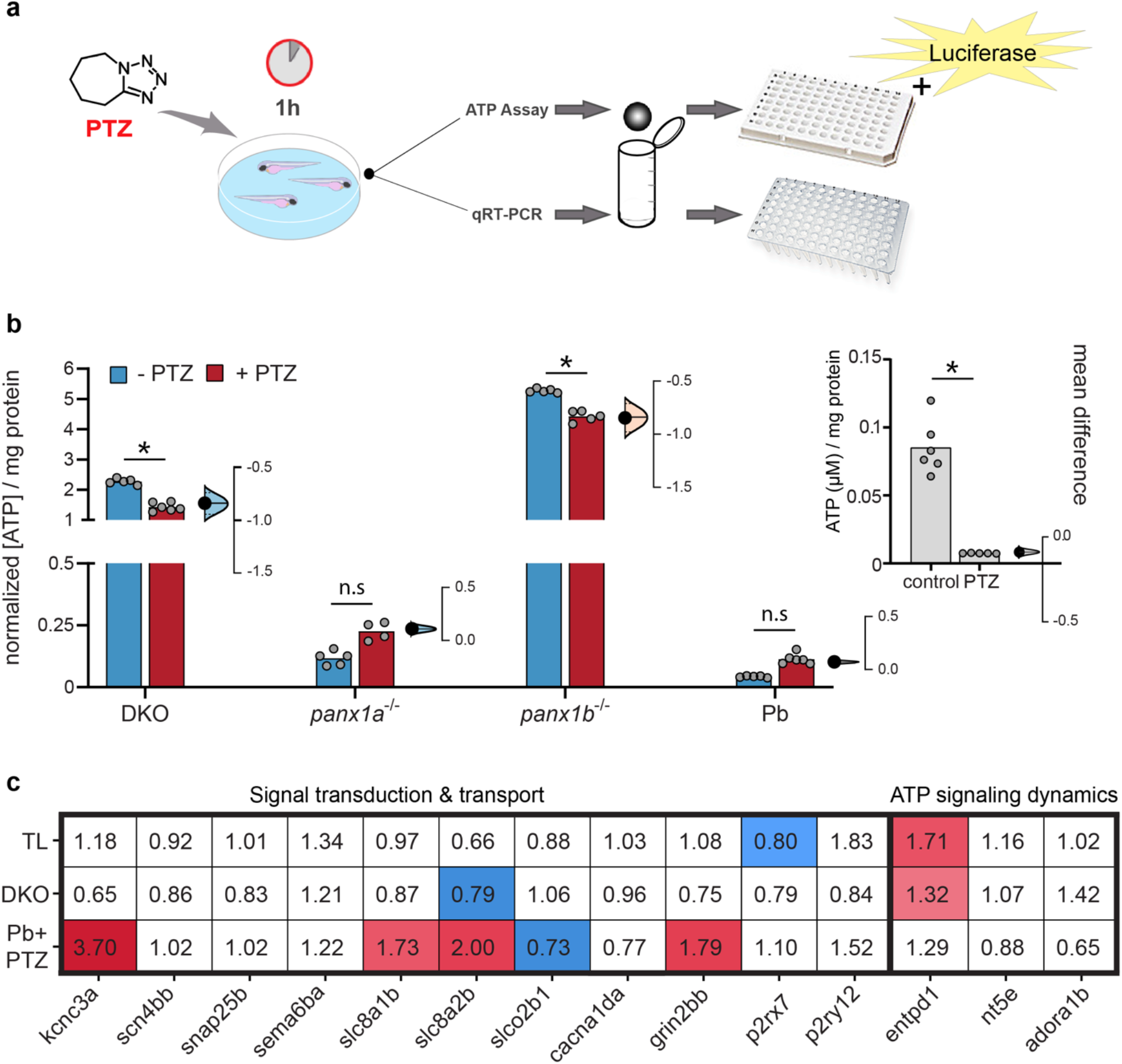
Extracellular ATP is associated with propensity for seizure-like activity. **a** Workflow diagram outlining the 1hr PTZ incubation for treated larvae. Larvae were collected for either RT-qPCR (n=30 larvae/sample) or ATP assays (n=50 larvae/sample), followed by homogenization prior to samples being plated in 96well format for measurement. Note for ATP detection, luciferase was used. **b** Estimation plots of extracellular ATP concentrations (µM) with respect to larval protein content (mg/ml) and normalized to the respective TL control. [ATP]_ex_ for TL controls are shown on the right (**b**), showing a significant decrease in ATP with PTZ treatment. Baseline ATP values are depicted by the blue bars, PTZ treatment is in red, individual data points are displayed in grey to show data distribution. Difference of means are shown to the right of each group, with a 95% confidence interval to show effect size and significance. DKO (Δ mean = -0.84) and *panx1b^−/−^* (Δ mean = -0.84) show a significant decrease in ATP with PTZ treatment. *Panx1a^−/−^* (Δ mean = 0.1) and TL larvae with Pb (Δ mean = 0.07) show very low ATP concentrations, however, do show a slight increase in ATP with PTZ treatment that is not significant. *Panx1b^−/−^* have the highest presence of ATP. **c** RT-qPCR of selected genes grouped into signal transduction and transport, or ATP signaling dynamic categories, showing significant up (red) or down (blue) regulation, with respect to non-treated controls, in response to PTZ treatment for TL, DKO and TLs treated with Pb. *p < 0.0001.

Reduced SLEs in gene edited DKOs and pharmacological blocked TL were not simply correlated with low [ATP]_ex_. This unexpected difference was corroborated by the differential expression of selected biomarkers for signal transduction and transport (**Fig. 7c, Supplementary Table 5**). Pb treated TL larvae showed an upregulation of voltage sensitive potassium channels required for rapid repolarization of fast-firing brain neurons (*kcnc3a*), Ca^2+^/Na^+^ antiporters (*slc8a1b* & *slc8a2b*) and NMDA receptors involved in excitatory postsynaptic potentials (*grin2bb*). DKOs had a significant downregulation of *slc8a2b*, which is an important Ca^2+^/Na^+^ antiporter found in axons and at the post synapse that plays a large role in Ca^2+^ and Na^+^ signaling and cell communication. Furthermore, PTZ treatment did not highlight a significant impact on the expression of genes associated with extracellular ATP processing, signal transduction or transport. The regulation of *entpd1,* which encodes for the rate-limiting ATP/ADP-hydrolyzing ectoenzyme CD39 appeared notable, but the expression of *p2rx7, p2ry12, nt5e*, or *adora1b* were similar in DKOs and Pb-treated TL. The results advocate for an important role of *panx1a* in ATP metabolism and signaling causing genotype-specific outcomes of seizure dynamics and allude to alternative mechanisms for acute versus chronic *panx1* targeting.

### *P2rx7* affects seizure-like activities but does not supersede *panx1a*

The impact of P2rx7 on seizure activity in the presence of both *panx1* genes was tested in TL larvae. The treatment with A-438079 hydrochloride hydrate (100µM A43; n = 46) started 30min before PTZ application and did not alter the larvae’s activity alone. PTZ-induced hyperactivity was reduced in A43 treated TL larvae compared to the activity induced in PTZ treated TLs (grey dotted line; n = 36) (**Fig. 8a**). The AUC analysis corroborated that treatment with both A43 and PTZ significantly reduced the AUC when compared to PTZ treated larvae, with and without extracted baseline activity (**Fig. 8b**). A43 treatment also significantly reduced Stage III counts (n = 18; **Fig. 8c**), without influencing the onset and peak activity of stages II and III in the test period.

**Fig. 8.**
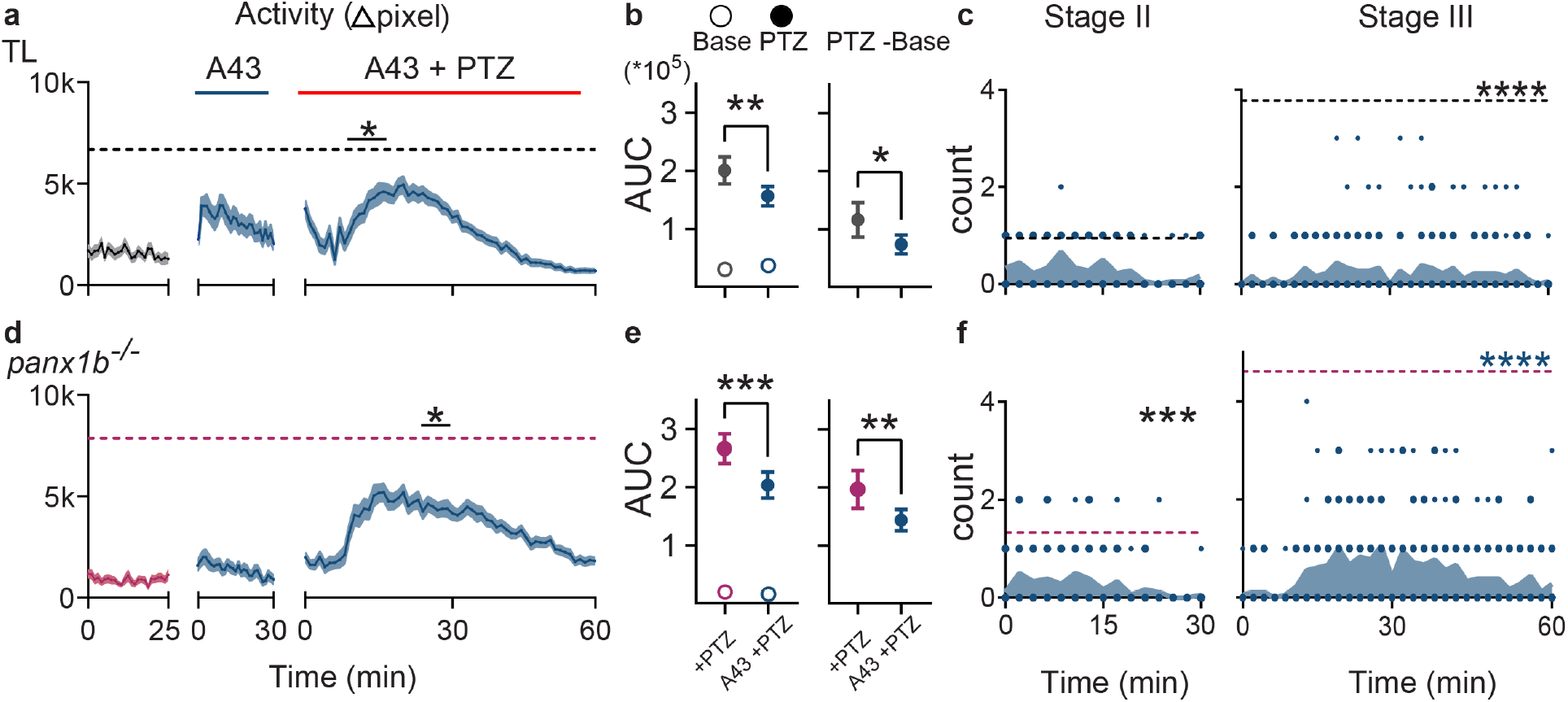
Blocking of P2rx7 with A-438079 improves seizure-like behavior. **a** 100µM A-438079 (A43) treatment decreased hyperactivity in TL in the first 10min of PTZ treatment (Δpixel ± s.e.m.; n = 46). Dashed grey line indicates TL’s max average activity score treated with only PTZ (n = 36). **b** AUCs for TL’s baseline activity and A43 treatment did not differ (left, open points). A43 treatment significantly reduced the effect of PTZ without (left, filled points) and with (right) extracted baseline activity. **c** Stage III count (count/2min) was significantly reduced with A43 treatment in TL (n = 18). Dashed lines represent max average Stage II and Stage III counts for TL’s PTZ-only group. **d** A43 treatment decreased PTZ-induced hyperactivity in *panx1b^−/−^* (n = 46). Dashed magenta line indicates *panx1b^−/−^*’s max average activity in the PTZ only group (n = 31). **e** A43 treatment significantly reduced the effect of PTZ in *panx1b^−/−^* without (left, filled points) and with (right) extracted baseline activity. **f** Stage II and III counts were significantly reduced with A43 treatment in *panx1b^−/−^* (n = 18). Dashed lines represent max average Stage II and Stage III counts for *panx1b^−/−^* ’s PTZ-only group. *p < 0.05, **p < 0.01, ***p < 0.001, ****p < 0.0001.

To isolate a potential role of *panx1a* in seizure-like activity, *panx1b^−/−^* larvae which express *panx1a* and *p2rx7* mRNA were treated with A43 (n = 46). The A43 treatment alone did not affect the baseline locomotor activity in *panx1b^−/−^* larvae. However, PTZ-induced activity was significantly reduced after 20min compared to activity in larvae treated with only PTZ (magenta dotted line; n = 31) (**Fig. 8d**). The AUC analysis confirmed that A43 treated *panx1b^−/−^* larvae had a reduced AUC compared to the PTZ treated, with and without extracted baseline activity (**Fig. 8e**). A43 treatment also reduced Stage II and III event counts significantly (n = 18; **Fig. 8f**). Our results demonstrated that the pharmacological targeting of P2rx7 in both TL and *panx1b^−/−^* larvae improved outcomes of PTZ induced seizure-like swimming behavior. Furthermore, we concluded that *panx1a* has a predominant role in seizure-like activity.

Furthermore, *panx1a* appears to play a predominant role in seizure-like activity.

### Molecular modeling of fish and human pannexins

High resolution cryo-EM structural studies of human and frog pannexins have provided an invaluable framework towards understanding the function of these channels ^40–43^. One unanticipated outcome of this work was that pannexins form heptameric channels in contrast to the hexameric organization observed for connexins gap junctions and the octameric organization observed for innexin gap junctions. In each of the four pannexin structural studies, a carboxy-terminal region with known regulatory activity was not observed and thus, its structure-function relationship could not be determined. Furthermore, many mechanistic questions remain with respect to gating and selection of ions despite having a high-resolution view of the channel.

A sequence alignment, shown in **Fig. 9a**, served a starting point for a structural comparison between human PANX1 and the two zebrafish pannexins. A high degree of sequence identity is evident, especially throughout the amino acids that line the channel **(Fig. 9b).** Two pairs of disulfide bonded cysteines in an extracellular facing domain of the channel are conserved in accordance with their established role in channel function ^44^.

**Fig. 9.**
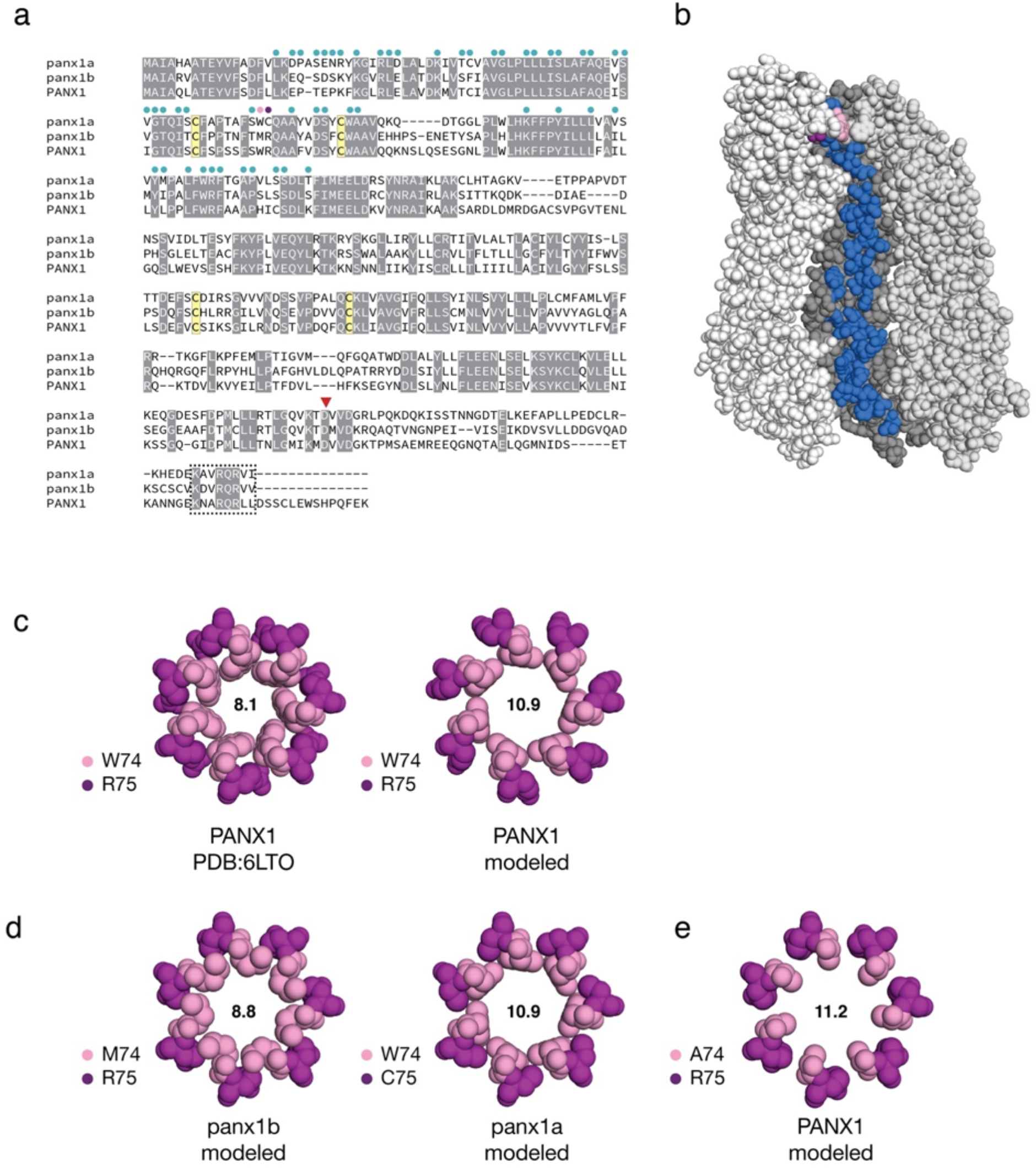
Sequence and structural attributes of the zebrafish pannexins. **a** Sequence alignment of human pannexin (PANX1) and zebrafish *panx1a/panx1b*. Amino acids that line the channel are indicated by circles above the sequence alignment. Two amino acids at positions 74 and 75 form an extracellular gate (in pink and purple, respectively). Four cysteines (in yellow) contribute two functionally important disulfide bonds. A caspase cleavage site is denoted by a red triangle. A box indicates a conserved carboxy-terminal segment of unknown significance. **b** Amino acids that line the inside of the channel are shown on one monomer of heptameric human PANX1 (PDB: 6LTO). The channel is oriented with the extracellular-facing side of the channel at the top. **c** The human PANX structure was used to model the extracellular gate of zebrafish *panx1a/panx1b* by making substitutions as required and repacking only amino acids 74/75 against a rigid backbone. As a control, human PANX1 was subjected to the same refinement and repacking protocol, creating an extracellular gate that was larger than the original cryo-EM structure. Diameters of the respective gates (in angstroms) are shown. **d** Molecular models of the extracellular gates of zebrafish *panx1a/panx1b*. Each protein bears one substitution relative to human PANX1. **e** An alanine was modeled to mimic a constitutively ATP permeable state. Molecular graphics were produced with PyMOL v2.4.1 (Schrödinger, LLC).

The pannexin channel has three gates each corresponding to a point of restriction **(Table 1)**. Among these gates, a sequence comparison of zebrafish Panx1a and Panx1b with human PANX1 revealed that the first and most narrow gate had enough sequence diversity to be explored further by molecular modeling. In human PANX1, the extracellular gate is described by W74 and R75. The guanidino group of R75 is positioned to support a favorable cation-pi interaction with the indole ring of W74 and an ionic bond with D81. Either or both interactions may be important for the function of the gate.

**Table 1:** A comparison of gates between human PANX1 and zebrafish Panx1a/Panx1b

|  | Human PANX1 | zebrafish Panx1a | zebrafish Panx1b |
| --- | --- | --- | --- |
| Gate 1 | W74-R75 | W74-C75 | M74-R75 |
| Gate 2 | I58 | V58 | V58 |
| Gate 3 | T21-E22-P23 | S21-E22-N23 | S21-D22-S23 |
| CBX* binding motif | I247-I258-F262 | I247-V258-L262 | L247-V258-V262 |

\*Carbenoxolone (CBX)

To produce models of the zebrafish Panx1a and Panx1b gates, the backbone of human PANX1 structure (PDB: 6LTO) was fixed in place, substitutions were made, and the side chains repacked. Before this was done, a control for the modeling study was performed by repacking the W74 and R75 side chains of human PANX1 according to the same protocol used to model the zebrafish Panx1a and Panx1b extracellular gates. The rotamer sampling method favored a wider gate due to an edge-to-edge packing of the indole ring of W74 (**Fig. 9c**) versus a staggered packing observed in the original cryo-EM structure increasing the diameter of the gate from 8.1 to 10.9 A.While the biological significance of these observed and modeled conformations are unknown, it demonstrates that side chain motions may limit the size of the extracellular facing gate in the absence of backbone conformational changes.

In zebrafish Panx1a, a cysteine (C75) replaces an arginine found in both human PANX1 and zebrafish Panx1b (**Fig. 9d**). While the loss of a positive charge and a nearby ionic bond could affect anion selection at this gate, a cysteine is still poised to support hydrophobic interactions with W74. In zebrafish Panx1b, methionine (M74) substitutes for tryptophan and the adjacent arginine (R75) is preserved. The modeled diameter of M74 was determined to 8.3A, comparable to W74 in the staggered conformation. To explain the reduced ATP transit that we have observed during our physiological experiments for Panx1b, we speculate that a methionine substitution may sample different conformations that either favor the closed state of the gate or simply create a gate that is smaller and thereby less permeable to ATP.

A W74A substitution is associated with increased channel activity, increased ATP release and a higher positive holding potential ^43^. When we modeled an A74 variant of human PANX1, the diameter of Gate 1 was 11.2 A (**Fig. 9e**). The distance is comparable to the 10.9 A observed in wild type PANX1 protein when the indole ring of W74 is in its most open edge-to-edge conformation. If the A74 variant extracellular facing gate is incapable of closing like the wild type protein, we question whether that is sufficient to explain the functional differences, or if there is an alternate ATP-permeable state that the A74 variant is incapable of forming. Towards answering this question, other amino acid substitutions at position 74 may be required.

## Discussion

The roles of Panx1 channels in neuronal excitability and experimental epilepsy models are controversial and incompletely understood ^45,46^. Here, the lower vertebrate zebrafish model was used to investigate pro-convulsant activities of *Panx1*. Our results demonstrate that a loss-of-function of the *panx1a* gene is the most discernable factor diminishing seizure susceptibility in gene-edited zebrafish by reducing extracellular ATP, and affecting biological processes related to transport and metabolism. In both gene-edited larvae and after acute blocking of *Panx1* channels, extracellular ATP and *P2rx7* were identified as factors contributing to seizure susceptibility. However, shared, and distinct outcomes of pharmacological and genetic models do not encourage a direct comparison of chronic and acute blocking of *Panx1* channels without caution.

To the best of our knowledge there is no direct evidence for Panx1 being epileptogenic and seizures are not a known comorbidity in patients with loss-of-function or gain-of-function mutations in human PANX1 ^47–49^. However, support for Panx1 contributing to seizures is from increased human PANX1 protein expression and seizure activity found in epileptic tissue ^8,11,12^. In rodents, activation of Panx1 augments aberrant bursting in the hippocampus and contributes to epileptiform seizure activities ^50^, meanwhile, inactivation leads to a reduction ^7–9,14^.

Here, blocking the gamma-aminobutyric acid (GABA)(A) receptor complex with PTZ evoked robust seizure-like events as reported previously in zebrafish ^22^ and mouse models ^7^. The two Panx1 ohnologues show both pro- and anti-convulsant activities in the presence of PTZ. Similar opposing activities in mouse Panx1 are cell-type and brain region specific ^15^. The genotype-specific differences described here are best explained by the distinct expression localizations and biophysical properties of Panx1a and Panx1b ^16–18^. Like mammalian Panx1 ^19–21^, Panx1a is broadly expressed throughout the zebrafish ^16,17^. However, the expression of Panx1b is more restricted to the nervous system ^17^. The biophysical properties vary when comparing the complexity of subconductance stages, the cumulative open and closed times, and the unitary conductance of panx1a (≈380pS) and panx1b (480-500pS) channels ^17^. These characteristics suggest that Panx1a channels copy characteristics of mammalian Panx1 and operate in different neuronal circuits of the brain to Panx1b.

Hypotheses regarding Panx1s’ involvement in epileptic seizures vary, with evidence pointing towards the ability to release the neurotransmitter ATP ^8,9 51^. Reports show extracellular ATP is reduced in brain tissue preparations deriving from mice with global loss of Panx1 ^9,52,53^, or astrocytic deletion of Panx1 ^14^, and extracellular ATP concentrations increase during high levels of neuronal activity and seizure-like events ^54,55^.

Since extracellular ATP levels depend on the rate of cellular release and enzymatic degradation, purinergic signaling, or the ratio between ATP and its metabolites ^54,55^, we quantified extracellular ATP and expression of genes involved in the process. In line with observations in mice, zebrafish lacking Panx1a channels show reduced ATP release efficiency, extending previously reported differences in channel activation kinetics and open times ^17,18^. Furthermore, RNA-seq analysis and a quantification of selected biomarkers provided evidence how ATP-related mechanism(s) contribute to pro-convulsant activities of Panx1a. The enriched biological processes and coordinated expression changes aligned with the opposite ATP release activities of *panx1a^−/−^* and *panx1b^−/−^* larvae, with DKO larvae representing mixed or bi-directional trends. Whether these opposing trends involve the differential expression of Panx1 channels in neurons and microglial cells known for protecting the mammalian brain from excessive activation that occurs during seizures ^56^ remains to be demonstrated. However, our results suggest that changes of the transcriptome represent a broad impact of Panx1 channels in different brain regions tuning the neuronal circuits they support based on their cellular expression and biophysical properties.

The recent description of Panx1 structures ^40,42,57^ allowed us to inquire the relationship of the zebrafish Panx1 ohnologues and the human paralog by structural modeling. Previous work by the Dahl group identified that ATP permeates ^58^ and activates mammalian Panx1 channels ^59^, and that amino acids W74 and R75 play a critical role in the process ^60^. Notable structural similarities and differences exist in the pore region of Panx1a, Panx1b, and human PANX1. However, the conservation of W74 at a critical gate in both human PANX1 and Panx1a suggests similar permeability of ATP in an open state,. We propose that this similarity enables Panx1a, like its mammalian paralogs to contribute to pro-convulsant activity via ATP signaling, as ATP release capabilities are reduced in the *panx1a^−/−^* larvae, after probenecid treatment, and in the DKOs. In turn, the lack of the conserved W74 could render Panx1b less competent for ATP signaling and seizure-like activities.

A functional interaction of the mammalian P2x7R-Panx1 complex is well established ^26^. Moreover, like Panx1, mammalian P2x7R expression is upregulated in seizure conditions ^61^. Although, targeting P2x7R in mice reduces epileptic seizures in some but not all models ^62–65^. Here, the downregulation of *p2rx7* in *panx1a^−/−^* larvae ^24^ coincides with a significant reduction of SLEs, suggesting that these channels work together. However, when P2rx7 receptors were blocked in TL and *panx1b^−/−^* larvae, only a moderate reduction of SLEs was observed, advocating for Panx1a and not P2rx7 as the principal driver of seizure-like activities.

In summary, the present study highlights the importance of *panx1a^−/−^* as the primary source for pro-convulsant activities in the zebrafish. We present models suitable for targeting different neuronal circuitries of the zebrafish, which will allow dissecting *in vivo* the mechanisms of how the brain is guarded from excessive excitability, and which functions require *Panx1*.

## Materials and Methods

### Zebrafish lines

All zebrafish (*Danio Rerio*) of strain Tupfel long fin (TL) were maintained in a recirculation system (Aquaneering Inc., San Diego, CA) at 28°C on a 14hr light/10hr dark cycle. All animal work was performed at York University’s zebrafish vivarium and in S2 biosafety laboratory in accordance with the Canadian Council for Animal Care guidelines after approval of the protocol by the Animal Care Committee (GZ: 2019-7-R2). The *panx1* mutant lines were generated in house ^24^.

### *In vivo* electrophysiology

Published procedures were used to prepare anaesthetized 7 days post fertilization (7dpf) zebrafish larvae for *in vivo* electrophysiology ^24,27^. Zebrafish larvae 7dpf were briefly anesthetized using 0.3mM Pancuronium bromide (Sigma-Aldrich) for 2-3 min until the touch response stopped. Anesthetized larvae were immobilized in freshly prepared 2% low melting temperature agarose. An Olympus dissecting microscope was used to orient the dorsal aspect of the larvae to the gel surface. Embedded larvae were placed on the upright stage of an Olympus BX51 fluorescence microscope. 1mL of egg water (E3; pH 7.2-7.4) was applied to the agar topically. Under direct visual guidance a glass microelectrode (1.2mM OD, approximately 1µM tip diameter, 2-7MΩ), backloaded with 2M NaCl, was placed into the right optic tectum. Local field potentials were recorded using a Multiclamp 700B amplifier (Axon Instruments, San Jose, CA, USA). Voltage recordings were low-pass filtered at 1kHz (-3 dB; eight-pole Bessel), high-pass filtered at 0.1 Hz, digitized 10 kHz using a Digidata 1550A A/D interface, and stored on a PC computer running pClamp11 software (all Axon Instruments). The basal activity was recorded for 10 minutes under Light-ON conditions (1000 lux), during which images of the electrode placement were taken for reference. Then, 1mL of E3 containing PTZ, for a final concentration of 15mM, was added topically to the agar and recorded for an hour. For each fish, seizure activity was normalized to its own baseline activity to account for biological variability of individual brains. For drug testing, zebrafish larvae were exposed to treatment drugs, 75µM Pb or 5mM VPA, at the start of baseline recordings 10min prior to the application of PTZ.

### Event detection and power spectral density

Ictal-like event identification was determined as high frequency events with large amplitudes (3 times the standard deviation of the baseline activity), polyspikes, and a duration of 3 seconds or greater. Inter-ictal-like events were identified as high-frequency events, with shorter amplitude compared to ictal-like events, but greater than baseline activity (minimum 1.5 times the standard deviation of baseline), and shorter in duration (1-3 seconds). Event detection was automated using custom developed codes in Matlab R2019b and visually confirmed ^28,29^. Events were quantified for comparison across genotypes and measured for significance using the Mann-Whitney test.

Power spectral density estimation was performed by the Welch’s method for baseline and PTZ recordings. A moving window for fast Fourier transform (FFT) computation was used and all windows were averaged. Changes in delta power were determined using the area under the power spectrum between 1-4Hz. The trapz function in Matlab R2019b was used for calculation. PSD was measured for significance using a one-way ANOVA (between genotypes) or an unpaired t-test (between treatments) and presented as the mean ± s.e.m..

### Zebrafish Locomotion Assays

The Zebrabox behavior system and the ZebraLab analytical suite (ViewPoint Life Technology, Lyon, France) were used for automated extraction of behavioral outputs and video tracking. Tracking videos were recorded at 30 frames per second (fps) under infrared light illumination using a Point Grey Research Dragonfly2 DR2-HIBW. A lightbox provided visible light for recordings at 30% light intensity. 7dpf larvae were observed in clear 96-well plates maintained at 28°C. Locomotor activity scores (Δpixel ± s.e.m.; n = 36 per group) were analyzed for larvae’s resting (25min), baseline (30min) and PTZ-induced behaviors (1hr). PTZ was administered in egg water at 15mM final concentration. In pharmacologically treated TL, Pb (75µM; n = 36), VPA (5mM; n = 60 per group), and A-438079 hydrochloride hydrate (A43; 100 µM; n = 46) were administered in egg water 30min prior to PTZ. Concentrations for Pb ^66^, VPA ^22^, and A-438079 ^67^ were chosen from the literature. A Pb dose-response test can be found here (n = 18; **Supplementary Fig. 4**). The Areas Under the Curve (AUCs) were determined from activity plots and analyzed by one-way ANOVA followed by a Tukey’s multiple comparison test, via an unpaired t-test or a Mann-Whitney test.

### Stage II and III seizure-associated behavior scoring

Videos extracted from the ZebraLab software were used for manually and blindly scoring stages II and III of seizure-associated locomotor behavior described in literature ^22^. Stage II manifests as ‘whirlpool-like’ rapid circling around the well (**Supplementary Video 1**) and stage III as a clonic-like convulsion followed by a loss of movement and posture (**Supplementary Video 2**). Stages II and III were scored at 2min intervals for a total of one hour after the PTZ application. Stage I, described as excessive hyperactivity, was not scored due to ambiguity and lack of quantifiable features for this stage, however, it is represented in the first 10min of the activity plots as a sharp increase in locomotor activity.

### Zebrafish larvae survival assessment

7dpf TL and *panx1* knockout larvae (n = 80 per group) were assessed for survival under brightfield microscopy every hour for a total of ten hours and once at 24-hours after the application of PTZ. Assessment included examining for circulation/heartbeat as well as body movement and degradation. Group survival rates were analyzed with the log-rank (Mantel-Cox) test and plotted as Kaplan Meier curves.

### Extracellular ATP assay

For quantification of ATP and protein ≈ 50 larvae 7dpf were collected at experimental endpoints. Larvae were homogenized in 500µl ice-cold of Phosphate-Buffered Saline (PBS) containing 100µM ARL-67156 (Sigma-Aldrich) and Halt Protease inhibitor (Thermo-Scientific) (1:100) for 1min at 30Hz using the Tissuelyser^LT^ (Qiagen). Homogenates were transferred to chilled Eppendorf tubes (^−^20°C) and centrifuged at 12,000 rpm for 2min. Supernatants were collected and snap-frozen in liquid nitrogen before storage at ^−^80°C. Extracellular ATP measurements used a 96-well format (Greiner Bio-One) and the Molecular Probes^®^ ATP determination Kit as described by the manufacturer (Life Technologies). ATP was quantified in replicates of 6 using the Synergy H4 Hybrid Multi-well Plate Reader (Biotek) as reported previously ^53^. Luminescent assay parameters; plate temperature set to 28°C; a low intensity shake of the plate for 3s prior to reading; a 5s integration time per well; gain setting at 150. In every test the ATP concentration in experimental samples was calculated from ATP standard curves (0-1µM ATP) included in the same 96-well plate. Data were exported from the Gen5 Data Analysis Software (Biotek) and analyzed in Excel and Matlab. The estimation plotting feature of Graphpad was used to test for statistical significance between treatment groups. ATP was represented as a normalized concentration per mg of protein. A NanoDrop Spectrophotometer was used to measure protein content in 2µL of supernatant from homogenized samples. The final ATP content was expressed with respect to the amount of protein content in each sample of pooled larvae after we had demonstrated that the weight of larvae and homogenate protein content were linearly correlated (**Supplementary Fig. 5**).

### RNA-seq and RT-qPCR

The transcriptomes of all zebrafish lines used in this research were analysed by RNA-seq (NGS-Facility, The Center for Applied Genomics, SickKids, Toronto, ON). The data derived from the sequencing of three independent pools of ≈30 age matched larvae (6dpf). Details of the bioinformatical analysis and the follow-up quantification of gene expression regulation by RT-qPCR can be found in Supplementary Information.

### Molecular modeling

The sequences of zebrafish Panx1a (Uniprot: Q7ZUN0) and Panx1b (Uniprot: F1QSR7) were aligned initially with Clustal Omega ^68^ to human PANX1 (Uniprot: Q96RD7). Using the alignment as a guide, zebrafish Panx1a and Panx1b were then threaded into the cryo-EM structure of human PANX1 (PDB: 6LTO) using SWISS-MODEL^69^. Using the FixBB module of Rosetta v3.12 ^70^, the side chains of amino acids 74 and 75 were altered and repacked following along the standard rotamer selection parameters of the program. Non-crystallographic symmetry (NCS) restraints were included to produce a heptameric assembly. The diameter of the extracellular gate was measured with PoreWalker ^71^.

### Pharmacology

All chemicals were purchased from Sigma-Aldrich (Mississauga, Canada): Pentylenetetrazole (PTZ; 15mM; cat#P6500), Pancuronium bromide (Panc; 300µM; cat#P1918), Probenecid (Pb; 75µM; cat#P8761), Valproic Acid (VPA; 5mM; cat# P4543) and A-438079 hydrochloride hydrate (A43; 100µM; cat#A9736). All concentrations referred to in the results are final.

### Statistics and reproducibility

All statistical analyses were performed in Matlab R2019b or in GraphPad Prism 9. Results are represented as the mean ± standard error of the mean (s.e.m.). For each analysis, a minimum of n ≥ 3 independent experimental replicates were generated. A *p-*value <0.05 was considered statistically significant. Unless otherwise stated, the Mann-Whitney test (nonparametric) and two-way repeated measures ANOVA (parametric) with the Greenhouse-Geisser correction followed by a Bonferroni’s multiple comparison test were used. G*Power analysis was used to confirm the required sample size and the statistical power of the acquired data^72^. For all experiments, sample sizes are indicated in the figure legends. Further statistical details are disclosed in **Supplementary Statistical Data.**

## Acknowledgements

We wish to acknowledge Uilki Tufa (University of Toronto, ON, Canada) for kindly assisting in developing automated seizure detection. Special thanks to Janet Fleites-Medina and Veronica Scavo for zebrafish husbandry. This research was supported by a Natural Sciences and Engineering Research Council (NSERC) discovery grants RGPIN-2018-05838 (LWD), RGPIN-2019-06378 (GRZ).

## Author contributions

Conceptualization, PW-F, GRZ; analysis, PW-F, DT, LCD.; investigation, PW-F, DT, NS, CZ; writing—original draft preparation, PW-F, DT, LCD; writing— review and editing, all authors; visualization, PW-F, DT, LCD; supervision, PLC, GRZ; project administration, GRZ; funding acquisition, GRZ. All authors have read and agreed to the published version of the manuscript.

## Materials & Correspondence

Correspondence and material requests should be addressed to Georg R. Zoidl

## References

1 Burnstock, G. Introduction to Purinergic Signalling in the Brain. Adv Exp Med Biol 1202, 1–12, doi:10.1007/978-3-030-30651-9_1 (2020).

2 Thompson, R. J. & Macvicar, B. A. Connexin and pannexin hemichannels of neurons and astrocytes. Channels (Austin) 2, 81–86, doi:10.4161/chan.2.2.6003 (2008).

3 Orellana, J. A. Physiological Functions of Glial Cell Hemichannels. Adv Exp Med Biol 949, 93–108, doi:10.1007/978-3-319-40764-7_5 (2016).

4 Ma, Z., Tanis, J. E., Taruno, A. & Foskett, J. K. Calcium homeostasis modulator (CALHM) ion channels. Pflugers Arch 468, 395–403, doi:10.1007/s00424-015-1757-6 (2016).

5 Mulligan, S. J. & MacVicar, B. A. VRACs CARVe a path for novel mechanisms of communication in the CNS. Sci STKE 2006, pe42, doi:10.1126/stke.3572006pe42 (2006).

6 Sabirov, R. Z. & Merzlyak, P. G. Plasmalemmal VDAC controversies and maxi-anion channel puzzle. Biochim Biophys Acta 1818, 1570–1580, doi:10.1016/j.bbamem.2011.09.024 (2012).

7 Aquilino, M. S. et al. Pannexin-1 Deficiency Decreases Epileptic Activity in Mice. Int J Mol Sci 21, doi:10.3390/ijms21207510 (2020).

8 Dossi, E. et al. Pannexin-1 channels contribute to seizure generation in human epileptic brain tissue and in a mouse model of epilepsy. Sci Transl Med 10, doi:10.1126/scitranslmed.aar3796 (2018).

9 Santiago, M. F. et al. Targeting pannexin1 improves seizure outcome. PLoS One 6, e25178, doi:10.1371/journal.pone.0025178 (2011).

10 Kim, J. E. & Kang, T. C. The P2X7 receptor-pannexin-1 complex decreases muscarinic acetylcholine receptor-mediated seizure susceptibility in mice. J Clin Invest 121, 2037–2047, doi:10.1172/JCI44818 (2011).

11 Jiang, T. et al. Altered expression of pannexin proteins in patients with temporal lobe epilepsy. Mol Med Rep 8, 1801–1806, doi:10.3892/mmr.2013.1739 (2013).

12 Li, S. et al. Expression of pannexin 1 and 2 in cortical lesions from intractable epilepsy patients with focal cortical dysplasia. Oncotarget 8, 6883–6895, doi:10.18632/oncotarget.14317 (2017).

13 Mylvaganam, S. et al. Hippocampal seizures alter the expression of the pannexin and connexin transcriptome. J Neurochem 112, 92–102, doi:10.1111/j.1471-4159.2009.06431.x (2010).

14 Scemes, E., Velisek, L. & Veliskova, J. Astrocyte and Neuronal Pannexin1 Contribute Distinctly to Seizures. ASN Neuro 11, 1759091419833502, doi:10.1177/1759091419833502 (2019).

15 Obot, P., Velisek, L., Veliskova, J. & Scemes, E. The Contribution of Astrocyte and Neuronal Panx1 to Seizures Is Model and Brain Region Dependent. ASN Neuro 13, 17590914211007273, doi:10.1177/17590914211007273 (2021).

16 Prochnow, N. et al. Pannexin1 in the outer retina of the zebrafish, Danio rerio. Neuroscience 162, 1039–1054, doi:10.1016/j.neuroscience.2009.04.064 (2009).

17 Kurtenbach, S. et al. Pannexin1 channel proteins in the zebrafish retina have shared and unique properties. PLoS One 8, e77722, doi:10.1371/journal.pone.0077722 (2013).

18 Bond, S. R., Wang, N., Leybaert, L. & Naus, C. C. Pannexin 1 ohnologs in the teleost lineage. J Membr Biol 245, 483–493, doi:10.1007/s00232-012-9497-4 (2012).

19 Vogt, A., Hormuzdi, S. G. & Monyer, H. Pannexin1 and Pannexin2 expression in the developing and mature rat brain. Brain Res Mol Brain Res 141, 113–120, doi:10.1016/j.molbrainres.2005.08.002 (2005).

20 Ray, A., Zoidl, G., Weickert, S., Wahle, P. & Dermietzel, R. Site-specific and developmental expression of pannexin1 in the mouse nervous system. Eur J Neurosci 21, 3277–3290, doi:10.1111/j.1460-9568.2005.04139.x (2005).

21 Baranova, A. et al. The mammalian pannexin family is homologous to the invertebrate innexin gap junction proteins. Genomics 83, 706–716, doi:10.1016/j.ygeno.2003.09.025 (2004).

22 Baraban, S. C., Taylor, M. R., Castro, P. A. & Baier, H. Pentylenetetrazole induced changes in zebrafish behavior, neural activity and c-fos expression. Neuroscience 131, 759–768, doi:10.1016/j.neuroscience.2004.11.031 (2005).

23 Grone, B. P. & Baraban, S. C. Animal models in epilepsy research: legacies and new directions. Nat Neurosci 18, 339–343, doi:10.1038/nn.3934 (2015).

24 Safarian, N., Whyte-Fagundes, P., Zoidl, C., Grigull, J. & Zoidl, G. Visuomotor deficiency in panx1a knockout zebrafish is linked to dopaminergic signaling. Sci Rep 10, 9538, doi:10.1038/s41598-020-66378-y (2020).

25 Cenedese, V. et al. Pannexin 1 Is Critically Involved in Feedback from Horizontal Cells to Cones. Front Mol Neurosci 10, 403, doi:10.3389/fnmol.2017.00403 (2017).

26 Iglesias, R. et al. P2X7 receptor-Pannexin1 complex: pharmacology and signaling. Am J Physiol Cell Physiol 295, C752–760, doi:10.1152/ajpcell.00228.2008 (2008).

27 Baraban, S. C. Forebrain electrophysiological recording in larval zebrafish. J Vis Exp, doi:10.3791/50104 (2013).

28 Hunyadi, B., Siekierska, A., Sourbron, J., Copmans, D. & de Witte, P. A. M. Automated analysis of brain activity for seizure detection in zebrafish models of epilepsy. J Neurosci Methods 287, 13–24, doi:10.1016/j.jneumeth.2017.05.024 (2017).

29 Colic, S., Wither, R. G., Zhang, L., Eubanks, J. H. & Bardakjian, B. L. Characterization of seizure-like events recorded in vivo in a mouse model of Rett syndrome. Neural Netw 46, 109–115, doi:10.1016/j.neunet.2013.05.002 (2013).

30 de Curtis, M., Jefferys, J. G. R. & Avoli, M. in Jasper’s Basic Mechanisms of the Epilepsies (eds th et al.) (2012).

31 Rakhade, S. N. et al. A common pattern of persistent gene activation in human neocortical epileptic foci. Ann Neurol 58, 736–747, doi:10.1002/ana.20633 (2005).

32 Losing, P. et al. SRF modulates seizure occurrence, activity induced gene transcription and hippocampal circuit reorganization in the mouse pilocarpine epilepsy model. Mol Brain 10, 30, doi:10.1186/s13041-017-0310-2 (2017).

33 Barkmeier, D. T. et al. Electrical, molecular and behavioral effects of interictal spiking in the rat. Neurobiol Dis 47, 92–101, doi:10.1016/j.nbd.2012.03.026 (2012).

34 Dahl, G., Qiu, F. & Wang, J. The bizarre pharmacology of the ATP release channel pannexin1. Neuropharmacology 75, 583–593, doi:10.1016/j.neuropharm.2013.02.019 (2013).

35 Turrini, L. et al. Optical mapping of neuronal activity during seizures in zebrafish. Sci Rep 7, 3025, doi:10.1038/s41598-017-03087-z (2017).

36 Safarian, N., Zoidl, C., Zoidl, G.. Pannexin1b is a candidate for setting interactions between the circadian clock and the zebrafish visual system. (2021).

37 Chen, E. Y. et al. Enrichr: interactive and collaborative HTML5 gene list enrichment analysis tool. BMC Bioinformatics 14, 128, doi:10.1186/1471-2105-14-128 (2013).

38 Kuleshov, M. V. et al. modEnrichr: a suite of gene set enrichment analysis tools for model organisms. Nucleic Acids Res 47, W183–W190, doi:10.1093/nar/gkz347 (2019).

39 Klepper, J. et al. Glut1 Deficiency Syndrome (Glut1DS): State of the art in 2020 and recommendations of the international Glut1DS study group. Epilepsia Open 5, 354–365, doi:10.1002/epi4.12414 (2020).

40 Deng, Z. et al. Cryo-EM structures of the ATP release channel pannexin 1. Nat Struct Mol Biol 27, 373–381, doi:10.1038/s41594-020-0401-0 (2020).

41 Mou, L. et al. Structural basis for gating mechanism of Pannexin 1 channel. Cell Res 30, 452–454, doi:10.1038/s41422-020-0313-x (2020).

42 Michalski, K. et al. The Cryo-EM structure of pannexin 1 reveals unique motifs for ion selection and inhibition. Elife 9, doi:10.7554/eLife.54670 (2020).

43 Qu, R. et al. Cryo-EM structure of human heptameric Pannexin 1 channel. Cell Res 30, 446–448, doi:10.1038/s41422-020-0298-5 (2020).

44 Bunse, S. et al. Single cysteines in the extracellular and transmembrane regions modulate pannexin 1 channel function. J Membr Biol 244, 21–33, doi:10.1007/s00232-011-9393-3 (2011).

45 Aquilino, M. S., Whyte-Fagundes, P., Zoidl, G. & Carlen, P. L. Pannexin-1 channels in epilepsy. Neurosci Lett 695, 71–75, doi:10.1016/j.neulet.2017.09.004 (2019).

46 Scemes, E. & Veliskova, J. Exciting and not so exciting roles of pannexins. Neurosci Lett 695, 25–31, doi:10.1016/j.neulet.2017.03.010 (2019).

47 Sang, Q. et al. A pannexin 1 channelopathy causes human oocyte death. Sci Transl Med 11, doi:10.1126/scitranslmed.aav8731 (2019).

48 Wang, W. et al. Homozygous variants in PANX1 cause human oocyte death and female infertility. Eur J Hum Genet, doi:10.1038/s41431-020-00807-4 (2021).

49 Shao, Q. et al. A Germline Variant in the PANX1 Gene Has Reduced Channel Function and Is Associated with Multisystem Dysfunction. J Biol Chem 291, 12432–12443, doi:10.1074/jbc.M116.717934 (2016).

50 Thompson, R. J. et al. Activation of pannexin-1 hemichannels augments aberrant bursting in the hippocampus. Science 322, 1555–1559, doi:10.1126/science.1165209 (2008).

51 Narahari, A. K. et al. ATP and large signaling metabolites flux through caspase-activated Pannexin 1 channels. Elife 10, doi:10.7554/eLife.64787 (2021).

52 Kurtenbach, S. et al. Investigation of olfactory function in a Panx1 knock out mouse model. Front Cell Neurosci 8, 266, doi:10.3389/fncel.2014.00266 (2014).

53 Whyte-Fagundes, P. et al. A Potential Compensatory Role of Panx3 in the VNO of a Panx1 Knock Out Mouse Model. Front Mol Neurosci 11, 135, doi:10.3389/fnmol.2018.00135 (2018).

54 Beamer, E., Conte, G. & Engel, T. ATP release during seizures - A critical evaluation of the evidence. Brain Res Bull 151, 65–73, doi:10.1016/j.brainresbull.2018.12.021 (2019).

55 Engel, T., Alves, M., Sheedy, C. & Henshall, D. C. ATPergic signalling during seizures and epilepsy. Neuropharmacology 104, 140–153, doi:10.1016/j.neuropharm.2015.11.001 (2016).

56 Badimon, A. et al. Negative feedback control of neuronal activity by microglia. Nature 586, 417–423, doi:10.1038/s41586-020-2777-8 (2020).

57 Ruan, Z., Orozco, I. J., Du, J. & Lu, W. Structures of human pannexin 1 reveal ion pathways and mechanism of gating. Nature 584, 646–651, doi:10.1038/s41586-020-2357-y (2020).

58 Bao, L., Locovei, S. & Dahl, G. Pannexin membrane channels are mechanosensitive conduits for ATP. FEBS Lett 572, 65–68, doi:10.1016/j.febslet.2004.07.009 (2004).

59 Locovei, S., Wang, J. & Dahl, G. Activation of pannexin 1 channels by ATP through P2Y receptors and by cytoplasmic calcium. FEBS Lett 580, 239–244, doi:10.1016/j.febslet.2005.12.004 (2006).

60 Qiu, F. & Dahl, G. A permeant regulating its permeation pore: inhibition of pannexin 1 channels by ATP. Am J Physiol Cell Physiol 296, C250–255, doi:10.1152/ajpcell.00433.2008 (2009).

61 Jimenez-Pacheco, A. et al. Increased neocortical expression of the P2X7 receptor after status epilepticus and anticonvulsant effect of P2X7 receptor antagonist A-438079. Epilepsia 54, 1551–1561, doi:10.1111/epi.12257 (2013).

62 Fischer, W. et al. Critical Evaluation of P2X7 Receptor Antagonists in Selected Seizure Models. PLoS One 11, e0156468, doi:10.1371/journal.pone.0156468 (2016).

63 Nieoczym, D., Socala, K. & Wlaz, P. Evaluation of the Anticonvulsant Effect of Brilliant Blue G, a Selective P2X7 Receptor Antagonist, in the iv PTZ-, Maximal Electroshock-, and 6 Hz-Induced Seizure Tests in Mice. Neurochem Res 42, 3114–3124, doi:10.1007/s11064-017-2348-z (2017).

64 Amhaoul, H. et al. P2X7 receptor antagonism reduces the severity of spontaneous seizures in a chronic model of temporal lobe epilepsy. Neuropharmacology 105, 175–185, doi:10.1016/j.neuropharm.2016.01.018 (2016).

65 Jimenez-Pacheco, A. et al. Transient P2X7 Receptor Antagonism Produces Lasting Reductions in Spontaneous Seizures and Gliosis in Experimental Temporal Lobe Epilepsy. J Neurosci 36, 5920–5932, doi:10.1523/JNEUROSCI.4009-15.2016 (2016).

66 de Marchi, F. O. et al. P2X7R and PANX-1 channel relevance in a zebrafish larvae copper-induced inflammation model. Comp Biochem Physiol C Toxicol Pharmacol 223, 62–70, doi:10.1016/j.cbpc.2019.05.012 (2019).

67 Donnelly-Roberts, D. L., Namovic, M. T., Han, P. & Jarvis, M. F. Mammalian P2X7 receptor pharmacology: comparison of recombinant mouse, rat and human P2X7 receptors. Br J Pharmacol 157, 1203–1214, doi:10.1111/j.1476-5381.2009.00233.x (2009).

68 Sievers, F. et al. Fast, scalable generation of high-quality protein multiple sequence alignments using Clustal Omega. Mol Syst Biol 7, 539, doi:10.1038/msb.2011.75 (2011).

69 Waterhouse, A. et al. SWISS-MODEL: homology modelling of protein structures and complexes. Nucleic Acids Res 46, W296–W303, doi:10.1093/nar/gky427 (2018).

70 Kuhlman, B. et al. Design of a novel globular protein fold with atomic-level accuracy. Science 302, 1364–1368, doi:10.1126/science.1089427 (2003).

71 Pellegrini-Calace, M., Maiwald, T. & Thornton, J. M. PoreWalker: a novel tool for the identification and characterization of channels in transmembrane proteins from their three-dimensional structure. PLoS Comput Biol 5, e1000440, doi:10.1371/journal.pcbi.1000440 (2009).

72 Faul, F., Erdfelder, E., Lang, A. G. & Buchner, A. G*Power 3: a flexible statistical power analysis program for the social, behavioral, and biomedical sciences. Behav Res Methods 39, 175–191, doi:10.3758/bf03193146 (2007).

