## Supplementary Information for "*Panx1* channels promote both anti- and pro-seizure-like activities in the zebrafish via *p2rx7* receptors and ATP signaling"

Supplementary Information (available in this document)

Supplementary Methods & References

Supplementary Figures 1–5

Supplementary Tables 1–6

Supplementary Videos 1,2

### Supplementary Methods

**TALEN Design -** Potential TALENs target sites on *panx1a* (NM_200916) and *panx1b* NM_001100030) were identified using Mojo Hand software (http://talendesign.org) [1,2]. The following criteria were used for TALEN design: TALENs target sites were 15-17 bases long with an initial 5' T nucleotide to the TALE domain. The spacer length was restricted to 15-16 base pairs. Target sites with a unique restriction enzyme sequence located in the middle of the spacer sequence were selected to simplify screening for insertion-deletion (indel) mutations. The specificity of selected TALENs target sequences was determined using the BLAST interface built into the Mojo Hand software.

**TALEN Constructs -** The TALEN constructs were synthesized in Dr. Stephen Ekker's lab (Mayo Clinic Cancer Center, Rochester, MN). Briefly, TALEN assemblies of the RVD-containing repeats were conducted using the Golden Gate approach [3]. Once assembled, the TALE repeats were cloned in the pT3TS-GoldyTALEN expression vector [4,5]. TALEN expression vectors were linearized with the *Sac*I restriction endonuclease (ThermoFisher Scientific, Canada) for 15 min at 37°C, and used as templates for *in vitro* transcription. Capped cRNAs were synthesized from TALEN pairs mixed 1:1 using the mMESSAGE mMACHINE T3 Transcription kit (Life Technologies, Canada). The mixture of the two TALEN cRNAs was purified using the Oligotex mRNA Mini Kit (Qiagen Inc., Toronto, Canada). TALEN cRNAs were diluted in DNase/RNase-free water (Life Technologies) to the final concentration of 1 μg/μL and stored at −80 °C before microinjection.

**Microinjection & genotyping -** One-cell stage zebrafish embryos were injected with TALEN cRNAs pair at doses ranging from 30-100 pg/nl. The toxicity of the injected cRNAs was determined at 24 hours post fertilization (1dpf) by calculating the proportion of healthy, dead, and malformed embryos at each dose. The condition resulting in more than 50% post-injection survival was selected for further injections. Genomic DNA (gDNA) was extracted from groups of 10 injected embryos at four days post fertilization (4dpf) to examine the TALEN mutagenesis efficiency. Briefly, the individual larva was incubated in 100mM NaOH at 95℃ for 15 min. After cooling to room temperature, the one-tenth volume of 1 M Tris (pH8.0) was added to the extracts to neutralize the NaOH [6]. Finally, 1 volume TE buffer pH8.0 was added, and gDNAs were stored at -20℃. A small indel mutation screen used PCR followed by *AfeI* (*panx1a*) and HindIII (*panx1b*) restriction enzyme (RE) digests. Indel mutations were confirmed by sequencing (Eurofins Genomics LLC, KY, USA) of gel-purified PCR products cloned into the pJet1.2 cloning vector (Life Technologies).

**Generation of knockout lines –** Adult mosaic zebrafish (F0) were anesthetized in pH-buffered 0.2mg/ml ethyl3-aminobenzoate methane sulfonate solution (MS-222, Sigma-Aldrich). The caudal fin (2 mm of the end) was removed using dissecting scissors (WPI Inc., FL, USA) and placed into 1.5 ml collecting tubes. The fin gDNA was isolated and screened for indel mutations as described [4]. Adult F0 zebrafish with desired indel mutations in the *panx1* genes were out-crossed to wild-type (WT) TL zebrafish. F1 offspring were analyzed by PCR and *AfeI/HindIII* digestions to verify germline transmission of mutations. Heterozygous F1 mutants were in-crossed to establish homozygous F2 mutants. Homozygous *panx1a*$^{-/-}$ and *panx1b*$^{-/-}$ lines were in-crossed to generate a double knockout line (DKO). All experiments described were performed with progenies of > F4 generations. Later generations were routinely tested for the identity of the genotype.

**RNA Extraction and RT-qPCR -** Total RNAs were extracted from pools of ≈30 7dpf larvae using RNeasy Plus Mini Kit (Qiagen). The iScript Reverse Transcription Supermix (Bio-Rad

Laboratories, Mississauga, Canada) was used to reverse transcribe 1µg of total RNA. The cDNA equivalent of ≈133ng total RNA was analyzed in triplicate by quantitative Real Time-PCR using the SsoAdvanced SybrGreen PCR mix (Bio-Rad). All experiments included a melt curve analysis of PCR amplicons generated in each reaction. Raw cycle threshold values (Ct-values) were exported from the CFX Manager Software (Bio-Rad, Canada), and the relative gene expression was calculated using the Relative Expression Software Tool (REST-2009) [7]. The statistical significance was tested by a Pair Wise Fixed Reallocation Randomisation Test©. Gene information and primer sequences are listed in **Supplementary Table 6**.

**RNA-seq Analysis** – The RNA library preparation was performed following the NEB NEBNext Ultra II Directional RNA Library Preparation protocol (New England Biolabs Inc., Ipswich, MA, USA). RNA libraries were loaded on a Bioanalyzer 2100 DNA High Sensitivity chip (Agilent Technologies) to check for size, quantified by qPCR using the Kapa Library Quantification Illumina/ABI Prism Kit protocol (KAPA Biosystems, Wilmington, MA, USA). Pooled libraries were paired-end sequenced on a High Throughput Run Mode flow cell with the V4 sequencing chemistry on an Illumina HiSeq 2500 platform (Illumina, Inc., San Diego, CA) following Illumina's recommended protocol to generate paired-end reads of 126-bases in length. The post-sequencing processing to final read counts, normalization, and differential gene expression analysis used multiple software packages, included RSEM version 1.3.3 (http://deweylab.github.io/RSEM/) and bowtie2 version bowtie/2.3.4.2 (http://bowtie-bio.sourceforge.net/index.shtml) to estimate the expression level of each sample. For each sample, RSEM reports read counts, estimated lengths and FPKM for each transcript and gene. For differential expression analysis estimated read counts for each transcript from RSEM output were compiled. This transcript expression matrix was supplied to DESeq2 (https://bioconductor.org/packages/release/bioc/html/DESeq2.html) v.1.22.2

to detect differentially expressed transcripts. Filtering of the low expressed transcripts to increase power was automatically applied via independent filtering on the mean of normalized counts within the DESeq results() function. Note: The transcripts that are filtered out have padj (FDR) value of "NA". The final output of the *DESeq2 results included* TranscriptID, GeneID – Ensembl IDs; GeneVersion, GeneName, GeneBiotype, baseMean - Mean of normalized counts for all samples; log2FoldChange - Log2 fold change*;* lfcSE - Standard error; stat - Wald statistic; *p*-value - Wald test *p*-value; padj - Benjamini+ Hochberg multiple testing; BH adjusted *p*-values.

**Transcriptome analysis -** Genes that were significantly regulated according to the padjusted value (<0.05) were organized by up (>1) and down (<-1) regulation based on the associated logFC value. Gene ontology (GO) enrichment of biological processes from the FishEnrichR database were established using these two gene lists for each genotype. Relevant biological processes, related to the nervous system and *panx1*, were categorized into broader GO terms for analysis and presentation. Clustergrams, were produced for differential expression analysis using curated gene lists and procedures implemented in the Matlab2019b Bioinformatics toolbox using Euclidean distance for hierarchal clustering.

### Supplementary Figures

**Supplementary Fig. 1** *Comparison of PTZ treatment to baseline activity is indistinguishable.*

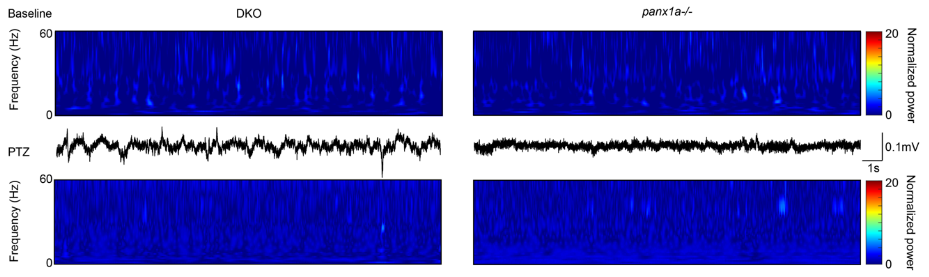

Spectrograms of baseline activity (top) directly compared to recordings during PTZ treatment (bottom) for DKO (left) and *panx1a-/-* (right) show that PTZ treatment does not affect neural activity in these genotypes. Sample traces from PTZ recordings at the 1hour mark are included for reference, revealing no major spiking activity.

**Supplementary Fig. 2** *Comparison of TL with and without Pb treatment only.*

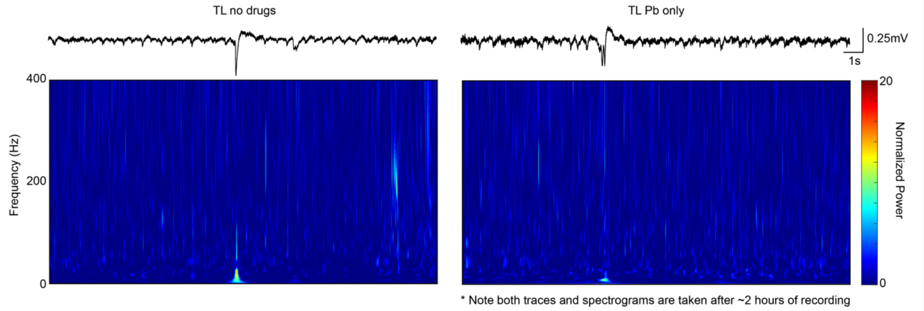

Spectrograms of baseline activity taken after 2hours of recording from TLs with no drug treatment (left) at all and TLs treated with probenecid only (right). This comparison shows that Pb is not inducing a toxicity effect to prevent seizure-like events from occurring as these spectrograms look like baseline activity.

**Supplementary Fig. 3** *Venn diagrams of differentially regulated genes and biological processes.*

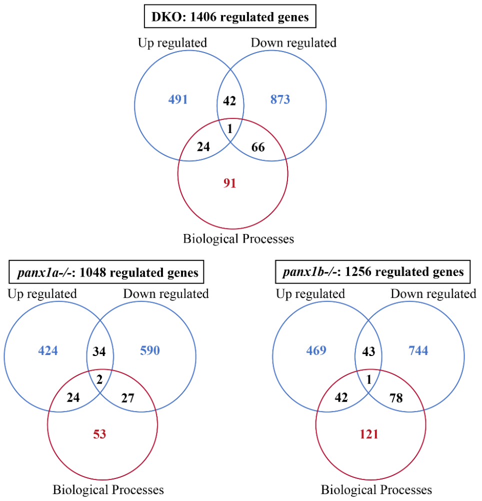

Venn diagrams highlighting how many genes are regulated in each of our *panx1* genotypes and show how many genes fit into biological processes according to FishEnrichR ontology database. They also reveal the overlap in regulated genes.

**Supplementary Fig. 4** *Concentration dependent reduction of PTZ-related hyperactivity in TL larvae by Probenecid.*

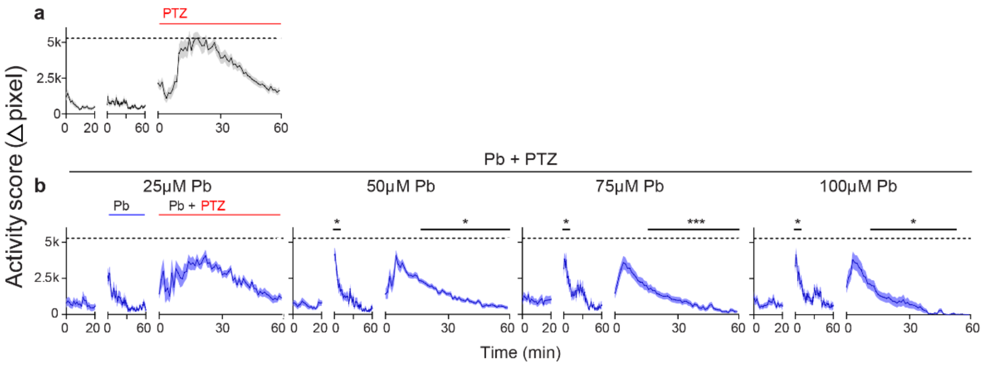

**a)** 7dpf TL larvae (n = 30) were treated with 15mM PTZ, or **b)** incubated with 25μM, 50μM, 75μM, and 100μM Probenecid (Pb; n = 18 per concentration) one hour before treatment with PTZ.

Pb significantly increased activity (Δpixel; mean ± s.e.m.) compared to baseline, and significantly reduced PTZ-induced activity above 50μM. Pb applied at 75μM reduced PTZ-induced activity, without the evidence of substantial toxicity (activity level 0 within one hour). Dashed lines indicate max average activity for PTZ treated TL. $*p < 0.05$, $***p < 0.001$.

**Supplementary Fig. 5** *Linear correlation of larval weight and protein concentration*.

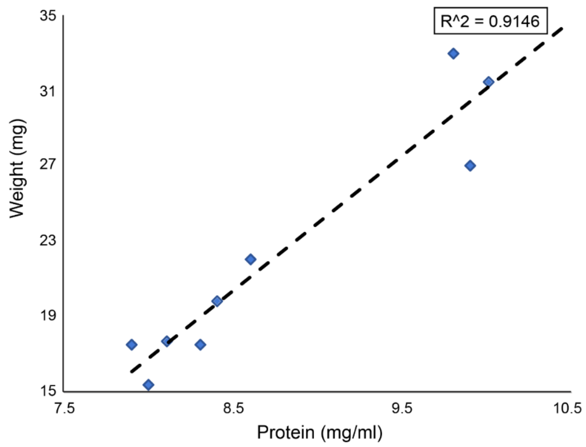

Plotting the weight of pooled larvae (n = 50 larvae/sample) in mg against the amount of protein in mg/ml that was measured from the supernatant of the homogenate with a spectrophotometer demonstrates a clear linear relationship ($R^2 = 0.9146$). Therefore, ATP concentrations were plotted per protein content to account for biological variance of larval weight.

### Supplementary Tables

**Supplementary Table 1** RT-qPCR values for Immediate Early Gene (IEG) regulation in TL and *panx1* knockout larvae treated with PTZ for one hour (**Fig. 4f**). Expression values, respective to non-treated controls, are >1 for upregulated IEGs and <1 for downregulated IEGs. *p*-values in bold = $p < 0.05$.

| Gene | TL | *p*-value | DKO | *p*-value | ***panx1a***$^{-/-}$ | *p*-value | ***panx1b***$^{-/-}$ | *p*-value |
|---|---|---|---|---|---|---|---|---|
| **fosab** | 110.36 | **0** | 67.62 | **0** | 88.89 | **0** | 57.72 | **0** |
| **egr1** | 9.16 | **0.001** | 7.35 | **0** | 10.41 | **0** | 6.30 | **0** |
| **egr2a** | 21.42 | **0** | 44.20 | **0** | 17.28 | **0** | 7.98 | **0** |
| **egr2b** | 6.86 | **0** | 4.65 | **0.002** | 7.59 | **0** | 5.32 | **0** |
| **egr4** | 120.54 | **0** | 63.68 | **0** | 76.99 | **0** | 44.53 | **0** |
| **jun** | 4.17 | **0** | 4.57 | **0** | 9.38 | **0** | 5.34 | **0** |
| **bdnf** | 6.13 | **0** | 3.95 | **0** | 4.73 | **0** | 3.10 | **0** |
| **eif4ebp2** | 1.04 | 0.765 | 0.96 | 0.813 | 1.56 | 0.05 | 1.50 | **0.01** |

**Supplementary Table 2** RT-qPCR values for IEG regulation in TL larvae treated with PTZ for one hour (+ PTZ) or treated with Pb 30min prior to the PTZ treatment (Pb + PTZ; **Fig. 5i**). Expression values, respective to controls not treated with PTZ (no treatment for + PTZ group and Pb treatment alone for Pb + PTZ group), are >1 for upregulated IEGs and <1 for downregulated IEGs. *p*-values in bold = $p < 0.05$.

| Gene | TL (+ PTZ) | *p*-value | TL (Pb + PTZ) | *p*-value |
|---|---|---|---|---|
| **fosab** | 110.36 | **0** | 54.61 | **0** |
| **egr1** | 9.16 | **0.001** | 6.09 | **0.001** |
| **egr2a** | 21.42 | **0** | 39.38 | **0** |
| **egr2b** | 6.86 | **0** | 2.03 | **0** |

| | | | | |
|---|---|---|---|---|
| **egr4** | 120.54 | **0** | 96.97 | **0** |
| **jun** | 4.17 | **0** | 1.44 | 0.073 |
| **bdnf** | 6.13 | **0** | 6.08 | **0** |
| **eif4ebp2** | 1.04 | 0.765 | 0.76 | 0.056 |

**Supplementary Table 3** Gene ontology enriched biological processes from FishEnrichR.

| DKO | Term from ZEBRAFISH ENRICHR with GO id | *p*-value | Log *p*-value | Z-score | Combined Score | Genes |
|---|---|---|---|---|---|---|
| **UP** | | | | | | |
| Transport | rRNA-containing ribonucleoprotein complex export from nucleus (GO:0071428) | 0.01 | 1.87 | -3.76 | 16.15 | npm1a;ran |
| | nuclear import (GO:0051170) | 0.05 | 1.31 | -2.02 | 6.10 | nup93;htatip2;ran |
| Signal transduction | regulation of calcium ion transmembrane transporter activity (GO:1901019) | 0.02 | 1.65 | -3.64 | 13.82 | cacnb4b;cacnb4a |
| | regulation of voltage-gated calcium channel activity (GO:1901385) | 0.04 | 1.41 | -3.84 | 12.42 | cacnb4b;cacnb4a |
| Metabolism | kynurenine metabolic process (GO:0070189) | 0.01 | 2.00 | -4.22 | 19.47 | afmid;tdo2a |
| | GTP metabolic process (GO:0046039) | 0.03 | 1.48 | -2.96 | 10.08 | ran;mbip |

| | | | | | | |
|---|---|---|---|---|---|---|
| Cellular respiration | - | | | | | |
| Cell death | neuron apoptotic process (GO:0051402) | 0.03 | 1.56 | -3.03 | 10.87 | pink1;siah1 |
| | mitochondrial outer membrane permeabilization (GO:0097345) | 0.02 | 1.75 | -3.19 | 12.80 | bnip3la;bnip3 |
| | positive regulation of mitochondrial membrane permeability involved in apoptotic process (GO:1902110) | 0.02 | 1.75 | -3.06 | 12.31 | bnip3la;bnip3 |
| **DOWN** | | | | | | |
| Transport | endosome transport via multivesicular body sorting pathway (GO:0032509) | 0.01 | 1.92 | -2.91 | 12.88 | tmem50a;vcp;chmp3 |
| | protein import (GO:0017038) | 0.05 | 1.33 | -2.14 | 6.58 | nup155;ipo7;ran;nutf2l |
| | protein localization to nucleus (GO:0034504) | 0.05 | 1.33 | -2.14 | 6.57 | nup155;ipo7;ran;nutf2l |
| Signal transduction | paraxial mesoderm development (GO:0048339) | 0.05 | 1.31 | -4.33 | 13.10 | kmt2a;gadd45aa |
| Metabolism | - | | | | | |
| Cellular respiration | regulation of reactive oxygen species metabolic process (GO:2000377) | 0.03 | 1.56 | -3.31 | 11.87 | mpx;acod1 |
| Cell death | | | | | | |

| | | | | | | |
|---|---|---|---|---|---|---|
| | intrinsic apoptotic signaling pathway by p53 class mediator (GO:0072332) | 0.01 | 1.92 | -3.64 | 16.12 | rps19;tp53;bag6l |
| | intrinsic apoptotic signaling pathway in response to endoplasmic reticulum stress (GO:0070059) | 0.03 | 1.56 | -7.08 | 25.40 | bag6l;tp63 |
| | intrinsic apoptotic signaling pathway in response to DNA damage by p53 class mediator (GO:0042771) | 0.04 | 1.42 | -5.65 | 18.54 | tp53;bag6l |
| ***panx1a-/-*** | **Term from ZEBRAFISH ENRICHR with GO id** | ***p*-value** | **Log *p*-value** | **Z-score** | **Combined Score** | **Genes** |
| **UP** | | | | | | |
| Transport | vesicle-mediated transport to the plasma membrane (GO:0098876) | 0.02 | 1.77 | -2.29 | 9.34 | stxbp6l;arf2a;exoc1 |
| | Golgi to plasma membrane transport (GO:0006893) | 0.03 | 1.51 | -1.18 | 4.10 | stxbp6l;arf2a;exoc1 |
| | post-Golgi vesicle-mediated transport (GO:0006892) | 0.04 | 1.37 | -1.56 | 4.94 | stxbp6l;arf2a;exoc1 |
| Signal transduction | response to axon injury (GO:0048678) | 0.05 | 1.35 | -2.37 | 7.33 | cnp;lingo1a |
| | axon regeneration (GO:0031103) | 0.05 | 1.29 | -2.55 | 7.60 | cnp;lingo1a |
| Metabolism | glutamine family amino acid catabolic process (GO:0009065) | 0.03 | 1.53 | -2.34 | 8.24 | aspg;glsb |
| | nucleobase-containing compound catabolic process (GO:0034655) | 0.05 | 1.35 | -1.78 | 5.51 | dnase1;rnaset2 |
| | cellular macromolecule | 0.06 | 1.25 | -1.79 | 5.13 | dnase1;rnaset2 |

| | | | | | | |
|---|---|---|---|---|---|---|
| | catabolic process (GO:0044265) | | | | | |
| Cellular respiration | cellular response to oxidative stress (GO:0034599) | 0.06 | 1.25 | -2.18 | 6.25 | pxmp2;prdx6 |
| Cell death | neuron apoptotic process (GO:0051402) | 0.02 | 1.68 | -3.04 | 11.76 | pink1;siah1 |

225

| | | | | | | |
|---|---|---|---|---|---|---|
| **DOWN** | | | | | | |
| Transport | nuclear transport (GO:0051169) | 0.01 | 1.87 | -4.29 | 18.50 | nxt2;nutf2l |
| | protein import (GO:0017038) | 0.01 | 1.87 | -2.19 | 9.43 | ipo11;pttg1ipb; nxt2;nutf2l |
| | protein localization to nucleus (GO:0034504) | 0.01 | 1.87 | -2.19 | 9.41 | ipo11;pttg1ipb; nxt2;nutf2l |
| | protein import into nucleus (GO:0006606) | 0.05 | 1.32 | -1.89 | 5.76 | ipo11;pttg1ipb; nxt2;nutf2l |
| Signal transduction | paraxial mesoderm development (GO:0048339) | 0.02 | 1.62 | -4.38 | 16.35 | kmt2a;gadd45a a |
| Metabolism | monocarboxylic acid catabolic process (GO:0072329) | 0.01 | 1.98 | -2.29 | 10.48 | acot8;aspg;acot 7 |
| | fatty acid catabolic process (GO:0009062) | 0.02 | 1.82 | -2.69 | 11.28 | acot8;acot7;aca a2;zgc:174917 |
| Cellular respiration | - | | | | | |
| Cell death | activation of cysteine-type endopeptidase activity involved in apoptotic process (GO:0006919) | 0.02 | 1.76 | -2.34 | 9.50 | hip1;diabloa;ca spbl |

| ***panx1b-/-*** | **Term from ZEBRAFISH ENRICHR with GO id** | ***p*-value** | **Log *p*-value** | **Z-score** | **Combined Score** | **Genes** |
|---|---|---|---|---|---|---|
| **UP** | | | | | | |
| Transport | cytoskeleton-dependent intracellular transport (GO:0030705) | 0.01 | 1.93 | -2.59 | 11.48 | kif1aa;ccdc88b;dync1i2a |
| | response to salt stress (GO:0009651) | 0.02 | 1.68 | -3.68 | 14.24 | slc26a6;trpv6 |
| Signal transduction | basement membrane organization (GO:0071711) | 0.02 | 1.68 | -3.80 | 14.73 | col4a6;pxdn |
| | regulation of calcium ion transmembrane transporter activity (GO:1901019) | 0.02 | 1.68 | -3.64 | 14.10 | cacnb4b;cacnb4a |
| | cytoplasmic microtubule organization (GO:0031122) | 0.03 | 1.51 | -2.03 | 7.07 | camsap2b;ccdc88b;fignl1 |
| | regulation of voltage-gated calcium channel activity (GO:1901385) | 0.04 | 1.44 | -3.83 | 12.70 | cacnb4b;cacnb4a |
| | regulation of cation channel activity (GO:2001257) | 0.05 | 1.32 | -2.29 | 6.99 | cacnb4b;cacnb4a;shank3a |
| | divalent inorganic cation homeostasis (GO:0072507) | 0.05 | 1.26 | -3.23 | 9.36 | trpv6;cnnm2b |
| Metabolism | GTP metabolic process (GO:0046039) | 0.03 | 1.51 | -2.96 | 10.30 | ran;mbip |
| Cellular respiration | - | | | | | |
| Cell death | | | | | | |

| | | | | | | |
|---|---|---|---|---|---|---|
| | - | | | | | |
| **DOWN** | | | | | | |
| Transport | nuclear import (GO:0051170) | 0.01 | 2.17 | -2.04 | 10.17 | pttg1ipb;htatip2;ran;nup62l;nutf2l |
| | rRNA-containing ribonucleoprotein complex export from nucleus (GO:0071428) | 0.03 | 1.55 | -3.71 | 13.19 | abce1;ran |
| | protein import into nucleus (GO:0006606) | 0.03 | 1.55 | -1.90 | 6.75 | abraa;pttg1ipb;ran;nup62l;nutf2l |
| | protein import (GO:0017038) | 0.03 | 1.54 | -2.16 | 7.65 | pttg1ipb;ran;nup62l;nutf2l |
| | protein localization to nucleus (GO:0034504) | 0.03 | 1.54 | -2.16 | 7.64 | pttg1ipb;ran;nup62l;nutf2l |
| | establishment of protein localization to organelle (GO:0072594) | 0.04 | 1.45 | -2.12 | 7.08 | pttg1ipb;ran;nup62l;nutf2l |
| | nucleocytoplasmic transport (GO:0006913) | 0.04 | 1.43 | -4.38 | 14.45 | htatip2;nutf2l |
| | ammonium transmembrane transport (GO:0072488) | 0.05 | 1.33 | -3.60 | 11.05 | rhcga;rhcgl1 |
| Signal transduction | calcium ion homeostasis (GO:0055074) | 0.05 | 1.31 | -1.59 | 4.79 | fhl1b;slc8a1a;atp2a2a;atp2b4;nr3c1 |
| Metabolism | negative regulation of cellular amide metabolic process (GO:0034249) | 0.01 | 2.16 | -2.53 | 12.58 | cnot7;fxr1;ormdl3;rpl13a |
| | gluconeogenesis (GO:0006094) | 0.02 | 1.80 | -1.99 | 8.22 | fbp2;tpi1b;pck1 |
| | glucose metabolic process (GO:0006006) | 0.05 | 1.32 | -2.22 | 6.75 | fbp2;tpi1b;pck1 |
| | cellular response to ketone (GO:1901655) | 0.04 | 1.43 | -3.71 | 12.25 | hamp;pck1 |
| Cellular respiration | | | | | | |

| | | | | | | |
|---|---|---|---|---|---|---|
| | regulation of reactive oxygen species metabolic process (GO:2000377) | 0.02 | 1.68 | -3.33 | 12.87 | mpx;acod1 |
| | mitochondrial electron transport, ubiquinol to cytochrome c (GO:0006122) | 0.05 | 1.33 | -3.58 | 11.01 | uqcrb;uqcrh |
| Cell death | - | | | | | |

**Supplementary Table 4** Clustergram genes and associated regulation values.

| Transport | | | |
|---|---|---|---|
| **Gene Name** | **DKO logFC value** | ***panx1a-/-* logFC value** | ***panx1b-/-* logFC value** |
| *exoc3l2b* | -0.5841 | - | - |
| *flot1b* | -0.5227 | - | - |
| *fthl28* | -1.399 | - | - |
| *gba* | -0.3936 | - | - |
| *hmgn3* | 0.4717 | - | - |
| *hspa8* | -2.3201 | - | - |
| *kcnc3a* | -7.5647 | - | - |
| *kcnf1a* | -5.2609 | - | - |
| *kcnk1a* | -2.3377 | - | - |
| *kcnk5a* | -1.4174 | - | - |
| *kdelr2a* | -0.5574 | - | - |
| *ktn1* | -0.898 | - | - |
| *lgals3b* | -1.0593 | - | - |
| *loxl2a* | -1.1992 | - | -1.1197 |
| *lrrc6* | -6.2773 | - | - |
| *mical3a* | -0.5704 | - | - |
| *mfsd4aa* | - | - | 0.781 |
| *myo5b* | -3.1089 | - | - |
| *ndel1b* | -0.5713 | - | - |
| *ndufa10* | -0.497 | - | - |
| *nlgn4xb* | - | - | -4.0171 |
| *npm1a* | -0.3634 | - | - |
| *nr3c1* | -8.2252 | 0.2801 | 0.3388 |

| | | | |
|---|---|---|---|
| *nsfb* | -0.8964 | - | - |
| *nutf2l* | -1.613 | - | - |
| *oc90* | - | 1.1787 | - |
| *p2rx7* | - | -1.1265 | - |
| *pafah1b1a* | -0.4651 | - | - |
| *panx1a* | -2.4621 | -2.7253 | - |
| *panx3* | -5.3551 | - | - |
| *pck1* | - | - | -1.0024 |
| *pink1* | 1.191 | - | - |
| *pitpnaa* | -1.7653 | - | - |
| *rab11bb* | - | - | -0.7261 |
| *rab18a* | 0.4016 | - | - |
| *rab3ip* | -8.4318 | - | - |
| *rock2b* | - | 1.436 | - |
| *rrbp1b* | -6.3568 | - | - |
| *scn4bb* | -5.5241 | - | - |
| *sdc2* | -0.4648 | - | - |
| *sdha* | - | - | -0.2181 |
| *sdhdb* | - | - | -0.5928 |
| *selenbp1* | 0.4764 | - | - |
| *shtn1* | -0.6307 | - | - |
| *slc12a4* | - | 0.4625 | - |
| *slc14a2* | -2.8632 | - | - |
| *slc16a3* | - | - | -8.0358 |
| *slc22a13a* | - | 3.5978 | - |
| *slc22a2* | - | - | -0.835 |
| *slc25a55a* | - | - | 0.5457 |
| *slc26a3.1* | -3.0933 | - | - |
| *slc26a6* | - | - | 3.1939 |
| *slc2a1a* | -0.861 | -0.9445 | - |
| *slc2a1b* | -1.0263 | - | - |
| *slc2a2* | - | - | 0.5076 |
| *slc34a2b* | 1.2019 | 5.3332 | 6.3308 |
| *slc3a2b* | -0.6289 | - | -1.0244 |
| *slc40a1* | 0.4478 | - | - |
| *slc43a1a* | -0.7094 | - | - |
| *slc43a2b* | -1.5258 | - | - |
| *slc44a2* | -0.6071 | - | - |
| *slc4a1a* | -7.9603 | - | - |
| *slc5a8l* | - | - | -0.7408 |
| *slc8a1b* | - | 0.6127 | 1.0189 |

| | | | |
|---|---|---|---|
| *slc8a2b* | -7.8466 | - | - |
| *slc9a3r1a* | - | 0.3518 | 0.392 |
| *snap25b* | -7.059 | - | - |
| *stim1a* | -1.2066 | - | - |
| *stra6* | -0.5599 | - | - |
| *tcnbb* | 0.6342 | - | - |
| *timm13* | -0.5591 | - | - |
| *tmed9* | -0.5686 | - | - |
| *trpv6* | - | - | 1.7513 |
| *tvp23b* | - | -0.6821 | - |
| *txnrd3* | -0.4692 | - | - |
| *ucp2* | - | - | -0.7065 |
| *ucp3* | - | - | -0.8651 |
| *uqcrb* | - | -0.3278 | - |
| *uqcrfs1* | -0.5015 | -0.487 | - |
| *vcp* | -8.8063 | - | - |
| *vps39* | -1.4432 | - | - |
| *yy1a* | 0.2932 | - | - |
| *zgc:92912* | - | - | -1.0469 |
| **Extracellular ATP** | | | |
| **Release** | | | |
| **Gene Name** | **DKO logFC value** | ***panx1a-/-* logFC value** | ***panx1b-/-* logFC value** |
| *p2rx7* | -1.1265 | - | - |
| *panx1a* | -2.7253 | - | -2.4621 |
| *panx3* | - | - | -5.3551 |
| *panx3* | - | - | -5.3551 |
| **Degradation** | | | |
| *enpp5* | - | -6.9338936 | - |
| *enpp7.1* | - | - | -1.033231287 |
| *entpd1* | - | - | -1.076657524 |
| *entpd5a* | - | - | -1.1842079 |
| *entpd5b* | - | -0.636278332 | - |
| **Metabolism** | | | |
| **Respirasome** | | | |
| **Gene Name** | **DKO logFC value** | ***panx1a-/-* logFC value** | ***panx1b-/-* logFC value** |
| *higd1a* | - | - | -1.7325 |
| *ndufa10* | - | - | -0.497 |
| *sdha* | - | -0.2181 | - |
| *sdhb* | - | -0.5928 | - |

| | | | |
|---|---|---|---|
| *uqcrb* | -0.3278 | - | - |
| *uqcrfs1* | -0.487 | - | -0.5015 |
| *mrpl12* | - | - | -0.4597 |
| *ndufa10* | - | - | -0.497 |
| *sdha* | - | -0.2181 | - |
| *sdhdb* | - | -0.5928 | - |
| *ucp2* | - | -0.7065 | - |
| *ucp3* | - | -0.8651 | - |
| *uqcrb* | -0.3278 | - | - |
| *uqcrfs1* | -0.487 | - | -0.5015 |
| **Solute Carriers** | | | |
| **Gene Name** | **DKO logFC value** | ***panx1a-/-* logFC value** | ***panx1b-/-* logFC value** |
| *slc1a5* | - | -0.5517 | - |
| *slc2a1a* | -0.9445 | - | -0.861 |
| *slc2a15a* | - | - | -1.7738 |
| *slc3a2b* | - | -1.0244 | -0.6289 |
| *slc6a2* | - | - | -1.0484 |
| *slc8a1b* | 0.6127 | 1.0189 | - |
| *slc8a2b* | - | - | -7.8466 |
| *slc20a1a* | - | -0.7041 | - |
| *slc22a2* | - | -0.835 | - |
| *slc22a13a* | 3.5978 | - | - |
| *slc25a4* | - | -0.7764 | - |
| *slc25a29* | 0.9005 | - | - |
| *slc25a36a* | - | 0.5108 | - |
| *slc25a40* | - | -0.7115 | - |
| *slc35d1b* | -0.7702 | - | -0.755 |

**Kommentiert [DT1]:** Anything supposed to be here or just empty cell?

**Supplementary Table 5** RT-qPCR values for gene regulation in PTZ treated (one hour) TL, DKO and Pb pretreated TL larvae compared to their respective controls (**Fig. 7c**). Expression values are >1 for upregulated genes and <1 for downregulated genes. *p*-values in bold = $p < 0.05$.

| **Gene** | **TL** | *p*-value | **DKO** | *p*-value | **TL + Pb** | *p*-value |
|---|---|---|---|---|---|---|
| ***kcnc3a*** | 1.18 | 0.319 | 0.65 | 0.311 | 3.70 | **0.001** |

| | | | | | | |
|---|---|---|---|---|---|---|
| ***scn4bb*** | 0.92 | 0.632 | 0.86 | 0.128 | 1.02 | 0.936 |
| ***snap25b*** | 1.01 | 0.946 | 0.83 | 0.346 | 1.02 | 0.9 |
| ***sema6b*** | 1.34 | 0.204 | 1.21 | 0.461 | 1.22 | 0.277 |
| ***slc8a1b*** | 0.97 | 0.895 | 0.87 | 0.302 | 1.73 | **0** |
| ***slc8a2b*** | 0.66 | 0.159 | 0.79 | **0.029** | 2.00 | **0** |
| ***slco2b1*** | 0.88 | 0.268 | 1.06 | 0.505 | 0.73 | **0.008** |
| ***cacna1da*** | 1.03 | 0.859 | 0.96 | 0.582 | 0.77 | 0.091 |
| ***grin2bb*** | 1.08 | 0.803 | 0.75 | 0.091 | 1.79 | **0.02** |
| ***p2rx7*** | 0.80 | **0.004** | 0.79 | 0.378 | 1.10 | 0.645 |
| ***p2ry12*** | 1.83 | 0.187 | 0.84 | 0.526 | 1.52 | 0.12 |
| ***entpd1*** | 1.71 | **0** | 1.32 | **0.02** | 1.29 | 0.07 |
| ***nt5e*** | 1.16 | 0.184 | 1.07 | 0.628 | 0.88 | 0.499 |
| ***adora1b*** | 1.02 | 0.959 | 1.42 | 0.133 | 0.65 | 0.331 |

**Supplementary Table 6** Primers for RT-qPCR.

| Gene | Gene Accession Number | Forward Primer (5'-3') | Reverse Primer (5'-3') |
|---|---|---|---|
| *fosab* | NM_205569 | GTGAACGAAACAAGATGGCTG | TTTCATCCTCAAGCTGGTCAG |
| *egr1* | NM_131248 | TCAACATATCCCAGTGCCAAG | TGTGTCTGGATGGGTTTCTG |
| *egr2a* | NM_001328404 | CTTCTCCTGTGACTTCTGCG | GCTTTCTGTCCTTATGTCTCTGG |
| *egr2b* | NM_130997 | GATGCGGAGAGGTCTATCAAG | AGGAGTAGGATGGCGGAG |
| *egr4* | NM_001114453 | ACAGCACCTCAAAGACTACAG | ACGACAAGGTAAAAGACTGGAG |
| *jun* | NM_199987 | CACAAGGCTCTGAAACACAAC | TGATGCCAGTTTGAGAAGTCC |
| *bdnf* | NM_001308648 | ACAAGCGGCACTATAACTCG | ACTATCTGCCCCTCTTAATGG |
| *eif4ebp2* | NM_212803 | AGTGACGGGCAAGAACATC | GTTGTTCACGTAGGTTCCTCTTC |
| *kcnc3a* | NM_001195240 | CCATGATAGGGCCTGCTTC | AGAGATGTTATTGAGGCTGCG |
| *scn4bb* | NM_001077573 | ACCTATGCCAGCTGTATTGG | CGCTCACGGTAAATTTGCAC |
| *snap25b* | NM_131434 | TGAGAATTTGGAGCAGGTCG | TGTTGGAGTCAGCCATGTC |
| *sema6ba* | NM_001366315 | TGATGGAGGGCTGTTTGTG | CGTTTGCGTGTTTGGGATC |
| *slc8a1b* | NM_001039144 | GGAGGGACCAGTTTATTGAGG | GGCACGAAAGCAAAGAGAAC |
| *slc8a2b* | NM_001123284 | TCACCAATGACCAGACAACTC | TGCACTCAACTGACCTTCTG |
| *slco2b1* | NM_001037678 | AGATGGATTGGTGCTTGGTG | TTCTCAGTTGATGGCTCCAC |

| | | | |
|---|---|---|---|
| *cacna1da* | NM_203484 | GGATGAGAAGGATAATGCCGAG | GGGTTTGTGTTGCTGAAGATG |
| *grin2bb* | NM_001128337 | ATGAGGGACAGGGATAGAGG | AGGTTGGGATGAATGGGTTC |
| *p2rx7* | NM_198984 | GTGTCATTTGTGGACGAGGAC | CACTCAACAGAGTCTTCATGCTG |
| *p2ry12* | NM_001308557 | TCTTCGGTTTGATCAGCATCG | TCAGGATTACATTTGGGAGCG |
| *entpd1* | NM_001003545 | ACCTGACCAACATGATTCCG | GCTGTTTTAGTAAAGCGACGG |
| *nt5e* | NM_200932 | CAAACGGAAATGTGCTGGAG | GTCTGTCCCACTTGCTGAG |
| *adora1b* | NM_001128584 | GGAACAATTTACACAGCCTGC | ACGAGCATGAAAAGCAGAGG |
| *tuba1a (ref)* | AF029250 | GAGCGTCCTACTTACACCAAC | AGGGAAGTGGATACGAGGATAG |
| *actb2 (ref)* | NM_181601 | GCCCCTAGCACAATGAAGATC | GACTCATCGTACTCCTGCTTG |

**Supplementary Videos**

**Supplementary Video 1** PTZ-induced seizure-associated behavior stage II: rapid 'whirlpool-like' circling around the well.

'ZF_SZR_STII_circling_behavior.avi'

**Supplementary Video 2** PTZ-induced seizure-associated behavior stage III: convulsions, uncontrollable twitch of the body followed by a loss of posture and movement.

'ZF_SZR_STIII_convulsive_behavior.avi'

1 Ma, A. C., Chen, Y., Blackburn, P. R. & Ekker, S. C. TALEN-Mediated Mutagenesis and Genome Editing. *Methods Mol Biol* **1451**, 17-30, doi:10.1007/978-1-4939-3771-4_2 (2016).

2 Neff, K. L. *et al.* Mojo Hand, a TALEN design tool for genome editing applications. *BMC Bioinformatics* **14**, 1, doi:10.1186/1471-2105-14-1 (2013).

3 Cermak, T. *et al.* Efficient design and assembly of custom TALEN and other TAL effector-based constructs for DNA targeting. *Nucleic Acids Res* **39**, e82, doi:10.1093/nar/gkr218 (2011).

4 Bedell, V. M. *et al.* In vivo genome editing using a high-efficiency TALEN system. *Nature* **491**, 114-118, doi:10.1038/nature11537 (2012).

5 Ma, A. C., Lee, H. B., Clark, K. J. & Ekker, S. C. High efficiency In Vivo genome engineering with a simplified 15-RVD GoldyTALEN design. *PLoS One* **8**, e65259, doi:10.1371/journal.pone.0065259 (2013).

6 Meeker, N. D., Hutchinson, S. A., Ho, L. & Trede, N. S. Method for isolation of PCR-ready genomic DNA from zebrafish tissues. *Biotechniques* **43**, 610, 612, 614, doi:10.2144/000112619 (2007).

7 Pfaffl, M. W., Horgan, G. W. & Dempfle, L. Relative expression software tool (REST) for group-wise comparison and statistical analysis of relative expression results in real-time PCR. *Nucleic Acids Res* **30**, e36, doi:10.1093/nar/30.9.e36 (2002).
